# NBS-SNI, an extension of the Network-based statistic: Abnormal functional connections between important structural actors

**DOI:** 10.1101/2022.08.29.505749

**Authors:** Francis Normand, Mehul Gajwani, Daniel C. Côté, Antoine Allard

**Affiliations:** Centre de recherche CERVO, Québec (Québec), Canada G1J 2G3; Centre interdisciplinaire en modélisation mathématique, Université Laval, Québec (Québec), Canada G1V 0A6; The Turner Institute for Brain and Mental Health and Monash Biomedical Imaging, Monash University, Melbourne, Australia; Département de physique, de génie physique et d’optique, Université Laval, Québec (Québec), Canada G1V 0A6

**Keywords:** Brain disorders, Structural Connectivity, Functional Connectivity, Centrality mea-sures, Anatomical measurements

## Abstract

Elucidating the coupling between the structure and the function of the brain and its development across maturation has attracted a lot of interest in the field of network neuroscience in the last fifteen years. Mounting evidence support the hypothesis that the onset of certain brain disorders is linked with the interplay between the structural architecture of the brain and its functional processes, often accompanied with unusual connectivity features. This paper introduces a method called the Network-based statistic–simultaneous node investigation (NBS-SNI) that integrates both representations into a single framework, and identifies connectivity abnormalities in case-control studies. With this method, significance is given to the properties of the nodes, as well as to their connections. This approach builds on the well-established Network-based statistic (NBS) proposed in 2010. We uncover and identify the regimes in which NBS-SNI offers a gain in statistical resolution to identify a contrast of interest using synthetic data. We also apply our method on two real case-control studies, one consisting of individuals diagnosed with autism and the other consisting of individuals diagnosed with early-psychosis. Using NBS-SNI and node properties such as the closeness centrality and local information dimension, we found hypo and hyperconnected subnetworks and show that our method can offer a 9 percentage points gain in prediction power over the standard NBS.

**AUTHOR SUMMARY:** We propose an extension to the well-known Network-based statistic (NBS) dubbed NBS-SNI, where the extension SNI stands for simultaneous node investigation. The goal of this approach is to integrate nodal properties such as centrality measures into the statistical network-based framework of NBS to probe for abnormal connectivity between important nodes in case-control studies. We expose the regimes where NBS-SNI offers greater statistical resolution for identifying a contrast of interest using synthetic data and test the approach with a real autism-healthy dataset which contains both the structural (**DTI**) and functional (fMRI) brain networks of each individual. We also tested our approach on a second dataset of individuals diagnosed with early-psychosis. In the second case, our framework is supplemented by incorporating the anatomically derived measures of intrinsic curvature index and gray matter volume directly as a node property, rather than the structural networks, thereby illustrating the versatility of our approach.

## 1 INTRODUCTION

Shedding light on the heterogeneous structure-function relationships in the brain has been the aim of many studies over the past fifteen years (Baum et al., 2017, 2020; Honey, Thivierge, & Sporns, 2010; Park & Friston, 2013; Pessoa, 2014; Suárez, Markello, Betzel, & Misic, 2020; Witvliet et al., 2021). These endeavors stem from the well established hypothesis in neuroscience that functional activity in the brain should be somehow related to its underlying structural architecture; indeed, the structural connectome is what supports the vast collection of functional patterns available to the human brain (Sporns & Kötter, 2004).

Correlations between brain structure and brain function have previously been demonstrated to be altered in schizophrenia, autism, bipolar disorder, small vessel disease, and mild cognitive impairment (Cao et al., 2020; Cocchi et al., 2014; Rudie et al., 2013; Tay et al., 2023; van den Heuvel et al., 2013; Wang et al., 2018; Zhang et al., 2019). This structurefunction coupling has also been shown to be prominent during youth, which would make it a crucial period for proper cognitive maturation (Baum et al., 2020).

However, integrated frameworks for concurrently analyzing both structure and function remain limited. Prior work has used disparate methodological approaches for relating brain structure and function, including the rich-club coefficient (van den Heuvel et al., 2013), individual or group level SC-FC correlational analyses (Cao et al., 2020; Tay et al., 2023; Wang et al., 2018), or primary component analysis (Rudie et al., 2013). These approaches may disregard information shared between structural and functional representations as structural and functional properties are assessed separately, and then combined. This may also result in confounds when comparing studies, due to the different statistical measure or metrics used.

One popular tool for evaluating differences in edge properties between (case and control) groups is the Network-based statistic (Zalesky, Fornito, & Bullmore, 2010). This approach has previously shown group level differences in functional connectivity edge weights in disorders including schizophrenia (Zalesky et al., 2010), autism (Pascual-Belda, Díaz-Parra, & Moratal, 2018), bipolar disorder (Roberts et al., 2017), and Alzheimer’s disease (Zhan et al., 2016). However, one key weakness of the NBS is that it only assesses properties of edges, and only assesses one edge property at a time. We look to extend the previous work undertaken in this direction by creating a framework that combines information from multiple streams, incorporating differences in both **nodes** and **edges** between groups.

Here, we introduce a procedure called Network-based statistic–simultaneous node investigation (NBS-SNI) which assesses properties of both nodes and edges, allowing for simultaneous assessment of two representations of the brain (e.g. structural and functional networks). In typical case-control paradigms, NBS-SNI aims to identify differences between two groups (case/control) of brain networks through an examination of abnormal connectivity between the most important **actors** of these networks. It does so by integrating two network representations of the brain at once, in pursuance of a more comprehensive representation of the brain. Within the proposed NBS-SNI framework, two properties are computed in each network of the healthy and the condition groups (one for each edge and one for each node), and compared through the lens of statistical testing. Combining information from nodes and edges in this way may increase both pathophysiological understanding and **statistical power**.

In Section 2, we give an overview of the general framework of the method and discuss potential choices of node properties that could be employed with our approach. In Section 3, we test our method with synthetic data to identify the regimes where our method NBS-SNI offers a statistical advantage over NBS or a generic link-based method. By considering the principal actors of the structural networks, we show that, in certain regimes, our method can flush out false positives and increase the resolution at which significant edges (true positives) can be identified. Finally, in Section 4, we apply our method on two real case-control datasets and perform classification of brain networks using various node properties. As an illustration, in the first dataset, using brain networks derived from neuroimaging data of a autism-healthy study (Rudie et al., 2013), where both the structural and functional networks are available (*N* = 29 ASD, *N* = 19 typically developing; TD), we found that NBS-SNI offers an improvement of 9 percentage points in prediction performance (73%) compared to the standard NBS (64%). On the second dataset, comprised of early psychosis-healthy individuals (*N* = 75 EP, *N* = 45 typically developing; TD), we incorporated anatomically derived measures (intrinsic curvature index and gray matter volume) as node properties in our proposed framework. We found that NBS-SNI offers a 4.9 percentage points gain compared to NBS.

## 2 METHODS

### Overview of the method

NBS-SNI is an extension to the well established Network-based statistic (NBS; Zalesky et al., 2010), a general framework used to test specific hypotheses in typical case-control studies. In essence, the formulation of NBS-SNI we propose tries to identify the most important nodes, common to the healthy and condition brain networks, and probes for abnormalities (hyper or hypo) in the interconnection around these influential brain regions. This is different from the standard NBS, which considers the extent to which significant edges (from a t-test perspective) are interconnected to form a component. The notion of component, i.e. retaining only the significant edges that form a connected component, is relaxed with this approach, since an additional restriction comes from the consideration of significant nodes, again by means of a statistical test, in addition to significant edges. A node property is computed for every node in the brain networks. The choice of a relevant node property in the context of brain network analysis is at the liberty of the investigator; a few possibly relevant node measures for network neuroscience are presented in section 2.

It is hypothesized that such a set of edges where stronger connections are observed between *particular* nodes in one group versus the other could be associated with certain symptoms, or network pathologies. Therefore, the choice of an appropriate node measure can be guided by prior knowledge of the specific condition under investigation. The resulting set of connections identified by this approach can then be paired with the framework NBS-predict proposed by Serin, Zalesky, Matory, Walter, and Kruschwitz (2021) to yield a prediction of individuals’ classes in case-control studies. In this paper, the relevance (significance) of the subnetworks extracted from real data are assessed by the prediction performances that they yielded on individual brain networks through the use of NBS-predict framework (as opposed to significance being assessed by permutation testing, as it was originally done in Zalesky et al. 2010).

This approach allows the analysis of both structural and functional representations of individual brain networks at once, under a single framework. Owing to the statistical framework of the method, which compares individuals across two representations of brain networks, a perfect **node correspondence** is required across both representations.

### Brain network representations

Depending on the imaging modalities employed, the nodes representing the constituents of the brain network representation could be individual neurons (micro-scale), groups of neurons (meso-scale) or brain regions (large-scale). The interactions (or associations) between the nodes are represented by the edges of the network, and their nature depends on the imaging technique employed. A brain network of *n* nodes is represented by a *n × n* connectivity (or adjacency) matrix, where the entries represent the pairwise associations between every possible pair of nodes.

The real brain networks investigated in this work originate from both fMRI (functional) and DTI (structural) recordings (large-scale) for every individual; micro/meso-scale brain networks will not be discussed here. Interactions measured by DTI structural imaging are bidirectional (undirected) edges whose weight corresponds to the density of white matter fibers connecting two brain regions. Interactions measured by functional imaging are also weighted and bidirectional, but their weight represents a measure of temporal correlation between the recorded dynamic activities between each pairs of parcellated brain regions.

### Node properties

NBS-SNI requires the use of a node property, which should be chosen to capture relevant node features related to efficient communication, resilience, clustering, or to the symptoms associated with the specific condition under investigation. Some of these node properties could be the degree, or the betweenness/closeness centrality, as they are well suited for large-scale brain networks dataset, in which the correspondence between the parcellated brain regions (the nodes) is reliable and the volume of each parcellated region does not vary much across individuals (Simpson, Lyday, Hayasaka, Marsh, & Laurienti, 2013). Node features could also be anatomically derived measures, such as the cortical thickness, gray matter volume, curvature, etc. The chosen node property is computed for every node in each network of the healthy and the condition groups, and then compared through the lens of statistical testing. Owing to the central limit theorem, it is worth mentioning that due to the smaller number of node properties (*n*) compared to the number of edge weights (*n*(*n −* 1)/2), the normality assumption may not always be reasonable for the computed node properties. In such a case, non-parametric statistical tests might be considered, instead of the usual student’s t-test, which will be used for edge weights statistical testing. Before getting into the details of these statistical tests, we briefly describe the node properties used here, and explain their relation to the notions of efficiency and resilience in brain networks. We also want to highlight the fact that the node properties don’t necessarily have to be computed from the structural brain networks. They could also be computed directly from the functional networks themselves. Moreover, in the second real dataset that we test with our tool-box, we used anatomically derived node measures from T1 images as a node property.

### Closeness centrality

The closeness centrality *C*_*C*_(*v*) of a node *v* quantifies how *close* it is, on average, to all the other nodes in the network. It is defined as the inverse of the average shortest path length *ℓ*_*v*_ (Newman, 2018)

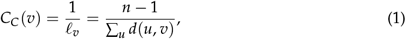

where *d*(*u, v*) is the shortest distance between *u* and *v*. Shortest distance here can be measured in terms of the smallest number of edges crossed to go from node *u* to node *v*, or in terms of the minimal “weight” accumulated while getting from nodes *u* to node *v*.

The interpretation of closeness centrality is usually one of access of efficiency or independence from potential control by intermediaries (Brandes, Borgatti, & Freeman, 2016). From this interpretation, a node of high closeness centrality would be less vulnerable to potential failures of other brain regions to which it is connected; thereby easing efficient rerouting of information along different pathways. In the context of case-control studies of brain networks, this interpretation of the closeness centrality makes it an interesting property to identify important nodes or identify between-group abnormalities. Closeness centrality will be used for both synthetic (see section 3) and real data (see section 4).

We also consider another measure called the current flow closeness centrality (sometimes called information centrality; Stephenson & Zelen, 1989). Contrary to the closeness, this node property does not make the assumption that communication happens solely along shortest paths (geodesics). Instead, information centrality uses all paths, and assigns them a weight based on the information they contain, therefore potentially capturing more subtle effects caused by non-geodesic paths in complex systems. This distinction with the closeness centrality makes it an interesting nodal property to investigate with NBS-SNI’s framework. The reader is referred to Stephenson and Zelen (1989) for detailed calculations of this measure. This measure will be used in section 4.

### Local Information Dimension

The last node property that will be investigated with NBS-SNI in section 4 with real data is called the local information dimension (LID) (Wen & Deng, 2020), a property derived from the local dimension (LD) Silva and da F. Costa (2013). For brain networks, a local definition of dimensionality allows one to quantify the local resilience of the network. The logic behind this statement is that objects of higher dimensionality might be less susceptible to being compromised by a random attack, since higher dimensionality often entails more _*redundant pathways, i*.*e*., *alternative information transmission pathways*._

With the LID, the number of nodes within a certain distance *l* of a central node *v* are considered using Shannon entropy. The size *l* represents the number of links starting from the central node *v* to the boundaries of the box. The LID 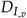 is defined as

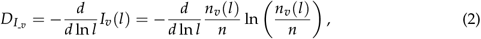

where 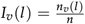 ln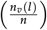 is the information contained in the box of size *l* whose central node is *v, n*_*v*_(*l*) is the number of nodes in a box of size *l* whose central node is *v*, while *n* is the total number of nodes in the network. With this measure, a larger value indicate a more “influential” node.

The LID 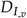 of a node *v* is obtained from the slope fitting of *I*_*v*_(*l*) versus ln *l*. In contrast to the LD, the size of the box *l* with LID varies from one to half of the maximum shortest distance from the central node *v*. This reduces the computational complexity and focuses on the *quasilocal* organization around the central node *v*. The LID was chosen over the LD as a node property to investigate in this paper owing to its reduced computational complexity, its greater effectiveness and reasonability at identifying influencers in complex networks (Wen & Deng, 2020). Additionally, we found that using the LID as a node property resulted in more accurate predictions than the (more global) local dimension (LD) (Silva & da F. Costa, 2013).

In section 4, we extend the LID to weighted networks by discretizing the *l* space into an arbitrary number of steps, starting from the minimum weighted shortest distance to half the maximum weighted shortest distance from the central node.

### Anatomically derived node measure

In the second real dataset that we investigated (see section 4), the anatomical measure of intrinsic curvature index (ICI) was employed as a node property. Its calculation was introduced by D. Van Essen and Drury 1997, and is defined as follows (Demirci & Holland, 2022):

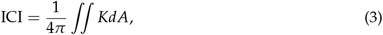

where *K* is the Gaussian curvature, defined as

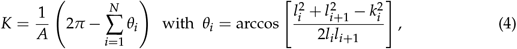

where *N* is the number of triangles connected to the vertex *p* (see Fig. 1). The measure of curvature is directly related to gyrification. Gyrification abnormalities have been reported in patients with schizophrenia (White, Andreasen, Nopoulos, & Magnotta, 2003; White & Hilgetag, 2011) and used to perform diagnosis of autism (Dekhil et al., 2019).

**Figure 1.**
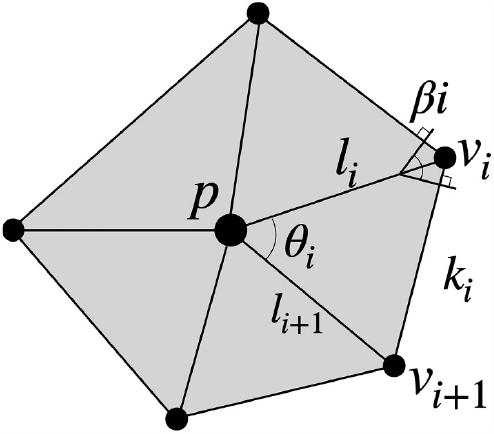
This figure and the following caption were taken directly from Demirci and Holland 2022. Representative polyhedra of the surface mesh at vertex *p* with *N* = 5 neighboring vertices. Edge *l*_*i*_ is associated with internal angle *θ*_*i*_ and dihedral angle *β*_*i*_ (note that vertex *p* is not necessarily co-planar with the neighboring vertices *v*_*i*_).

Finally, the gray matter volume (GMV) of each parcellated brain region was also employed as a node property with the HCP early psychosis dataset in section 4. Gray matter volume reductions and abnormalities have been reported in patients with schizophrenia (Gur et al., 2000; Gur, Turetsky, Bilker, & Gur, 1999; Hulshoff Pol et al., 2002; Vita, De Peri, Deste, & Sacchetti, 2012) and also in patients with autism (Dekhil et al., 2019), thereby making it a suitable node property to incorporate into NBS-SNI.

### NBS-SNI

In the following description, the method is designed to identify the most important nodes (brain regions) common to both the condition and the control group in the structural representation, and probe for hyperconnected (or hypoconnected) functional connections between these brain regions.

In what follows, both the structural and functional connectivity matrices are available for each individuals in both groups (case and control). Moreover, the method assumes that there is a “perfect” correspondence between the nodes across the networks for both representations (*node-aligned networks*). Nevertheless, we want to point out that it would also be reasonable to compute the node properties directly from the functional connectivity matrices. Furthermore, node properties could also be derived from anatomical measurements, without necessarily having to be extracted from the structural connectivity matrices. This will be illustrated in section 4, where the anatomically derived measures of intrinsic curvature index and gray matter volume will be used as a node property with NBS-SNI.

Let there be *N*_1_ pairs of networks (functional and structural) of *n* nodes in group 1 (condition) and *N*_2_ pairs of networks of *n* nodes in group 2 (control). To find significant nodes common to both the condition and control group, a node property is computed for each node from the structural connectivity matrices. This results in a *N*_1_ *× n* matrix where each row contains the node measures of all the nodes in a structural network from group 1. The same procedure is done with the other group to obtain a *N*_2_ *× n* matrix of node properties (step 1 on Fig. 2). Two independent one-sample statistical tests are then performed on this property, yielding two vectors of length *n* containing the statistic of each node, i.e., one vector for each group (step 2). It is important to note at this point that the normality assumption required to apply the t-test might not always be met when using node properties due to the small number of nodes (compared to the number of edges). In that case, the user might resort to a non-parametric test such as the Kolmogorov-Smirnov test instead of the one-sample t-test.

**Figure 2.**
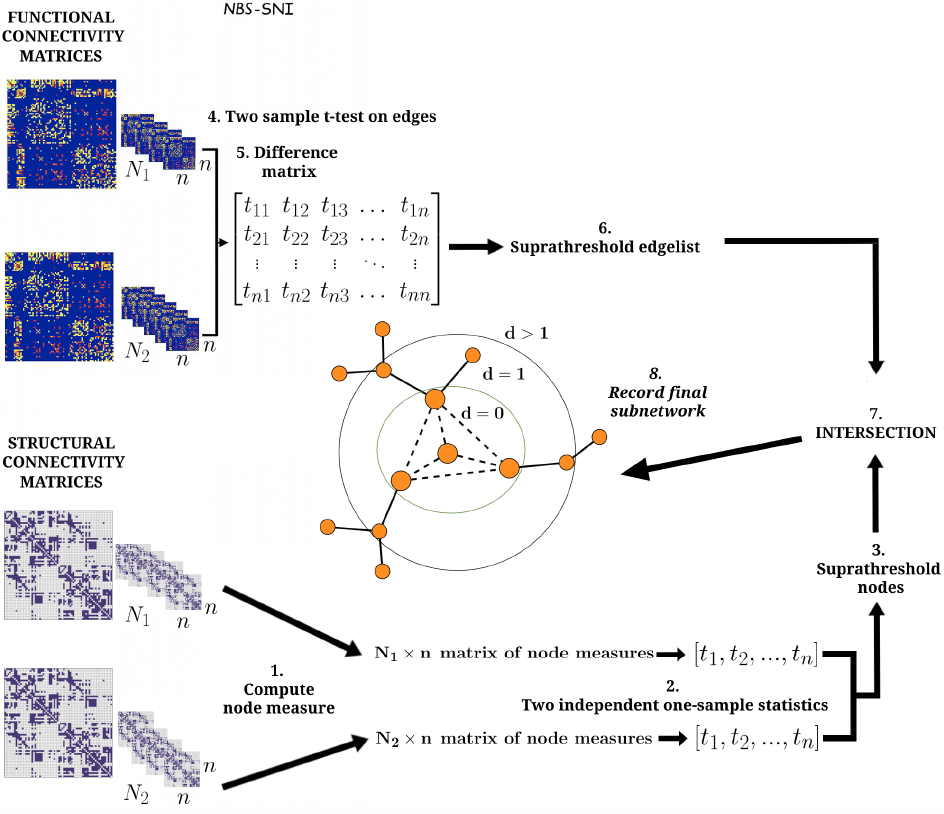
Illustration of NBS-SNI. The inputs are the structural and the functional connectivity matrices, two matrices for each individual in the condition and in the control groups. Statistical tests are performed in parallel on both sets of matrices (structural and functional). On the functional representation, a two-sample t-test is performed on the entries of the connectivity matrices (edge weights) (step **4**), which are then thresholded (step 5), yielding a set of suprathreshold edges (step 6). On the structural representation, a node measure is computed for every node (step **1**), and a one sample statistical test is performed on these node measures (step **2**) independently for each group. Both sets of nodes’ statistical scores are then thresholded. At this point, we suggest taking the intersection between the resulting suprathreshold nodes (step **3**), thus retaining only the nodes that were statistically significant in both the condition and the control group. Alternatively, the investigator could select only one supratheshold nodes’ set (for e.g., if one set contains many supratreshold nodes while the does not contains only a few, or if the overlap is empty). Finally, the intersection (*d* = 0) (or a relaxed intersection (*d≥*1)) (step **7**) between the suprathreshold nodes and the suprathreshold edges is taken, and the final set of edges remaining is recorded (step **8**). The three cases considered shown above in the center are *d* = 0, *d* = 1 and *d >* 1, where *d* is the distance of a node from the central nodes, which are depicted slightly larger.

A threshold *t*_node_ is then applied to the two resulting **statistics** independently in each group to identify significant nodes, and the intersection between the significant nodes of both groups is taken (step 3). These **suprathreshold** nodes correspond to the most important actors (nodes) of these networks mentioned earlier. Alternatively, it would be reasonable to retain only the suprathreshold nodes from one of the two groups if the intersection between the two sets is empty, or if one set contains many suprathreshold nodes, while the other contains a few.

In parallel, a two-sample t-test is also performed on the edge weights of the functional connectivity matrices for a between-group difference (step 4), resulting in a single *n × n* matrix that contains the t-statistic of each edge weight as its entries. A threshold is then applied to this matrix (step 5), resulting in a set of suprathreshold edges (step 6).

Finally, the intersection between the resulting suprathreshold nodes and suprathreshold edges is taken (steps 7 and 8). Three cases naturally arise. The most restrictive case consist in retaining only the edges that are shared between the “important” nodes (*d* = 0).

Note that for *d* = 0, the subnetwork yielded by NBS-SNI is always a connected component. The second case implies retaining all edges that connect at least one suprathreshold node (*d* = 1). The third case retains edges that connect nodes that are at most at a distance *d >* 1 from the suprathreshold nodes. Note that this third case requires computing shortest distances from the central nodes, which increases the computational complexity of the method. Hereinafter, we chose to perform the statistical test on the edge weights of the functional connectivity matrices, while the statistical test on the node properties takes place on the structural connectivity matrices, but swapping these choices could be valid as well. Also, note that an alternative formulation where the investigator first probes for abnormalities in the nodes from the structural representation could also be valid (i.e., performing a two-sample statistical test for between-group differences), as opposed to finding nodes considered important and common to both the condition and control group. This formulation employing a two-sample t-test will be used with a real dataset in section 4 using the anatomical measure of gray matter volume as a node property on the HCP early psychosis dataset.

In our simulations presented in section 3, NBS and NBS-SNI are sometimes compared against the false discovery rate (FDR), i.e. the expected ratio of false positives among all the positives, which serves as a link-based controlling procedure. With FDR, every edge’s t-score is treated independently, as opposed to NBS or NBS-SNI, which do not yield a collection of individual significant edges, but rather a connected component that they form.

### Classification framework

We adapted the framework NBS-predict (Serin et al., 2021) to investigate the impact information about nodes taken into account by NBS-SNI has on the accuracy of a classification task. In this context, individual brain networks (functional connectivity matrices) are classified (diagnosed) into either the control or the condition group. The edge weights of the subnetworks extracted by the method are the features which are fed to various machine learning (ML) algorithms (the features associated to the edges that were not selected are set to 0). A *K*-fold cross validation procedure is repeated *r* times (see Fig. 3). Finally, the method outputs an average prediction accuracy for the brain networks and a weighted adjacency matrix, in which the weights of the edges represent their individual prediction accuracy across folds. Figure 3 illustrates the workflow of the classification procedure NBS-predict.

**Figure 3.**
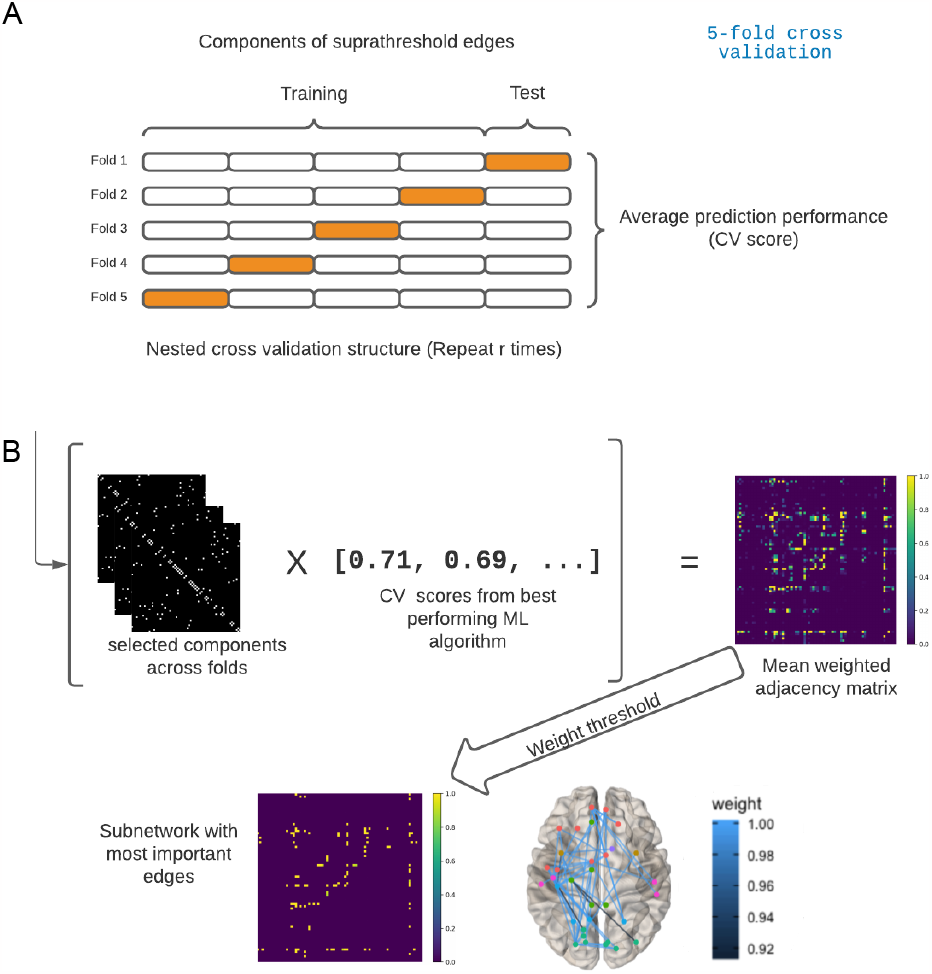
Illustration of the process employed for the classification task. A. Nested crossvalidation structure (*K* = 5 folds in this example) employed to yield out-of-sample predictions. A nested cross validation is one that is repeated *r* times. This is done to take into account possible variations in the prediction performances in the test subsets. B. The framework of NBS-predict (Serin et al., 2021). The subnetworks binary adjacency matrices obtained across folds are multiplied with their associated prediction performance. This procedure outputs a weighted adjacency matrix, which holds the relative importance of each edge in making predictions across the nested cross-validation process. It is then possible to apply a threshold to the resulting weighted adjacency matrix to retain and visualize only the edges that contributed the most to the predictions. This illustration is inspired by Fig 1 from Serin et al. (2021).

To visualize the contribution of each edge to the overall prediction performance, the selected edges in the extracted subnetworks across folds are constantly updated and assigned a weight (encoded in a single weighted adjacency matrix), which corresponds to the prediction performance at that specific fold (here again, edges that are not selected are assigned a weight of 0). In the end, all the weights are averaged and rescaled so that the maximum prediction weight is 1. One may then visualize the selected edges across folds that contributed the most to the prediction (i.e., the largest weights denote the edges that had a greater prediction accuracy and that were selected the most across folds).

The classification ML algorithms employed in this work are Ridge Classifier, Random Forest Classifier, Decision Tree, Support Vector Machine (SVM) and Logistic Regression, all implemented from the scikit-learn (Pedregosa et al., 2011) Python package.

## 3 CONTRAST IDENTIFICATION: NBS-SNI VS. NBS USING SYNTHETIC DATA

To compare the statistical resolution of NBS-SNI with NBS to identify a **contrast**, synthetic data was created for both structural and functional connectivity brain networks in a typical case-control study setup. As mentioned previously, we expect NBS-SNI to draw its strength from considering both representations simultaneously, when an interplay of contrasts exists across the two representations.

### Synthetic data model

#### Generative model for the structural networks

Our approach to generate synthetic structural networks is inspired by Akarca, Vértes, Bullmore, and Astle (2021); Betzel et al. (2016) and goes as follows. Edges are added one at a time probabilistically until the desired number of edges has been added. The probability of forming an edge between node *u* and node *v* is given by

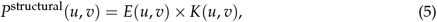

where *E*(*u, v*) represents the probability of connection between brain regions *u* and *v*, depending on the Euclidean distance *d*_*uv*_ separating them. The second term *K*(*u, v*) assigns a greater probability of connection on the preselected set of nodes, *n*_*c*_, defined as the contrasting nodes, or alternatively adds noise to the connection probability between the other nodes. These rules for *K*(*u, v*) are expressed as

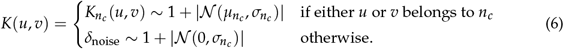

By increasing the connection probability between *u* and *v* if either belongs to *n*_*c*_, the last term in Eq. 5 effectively increases the closeness centrality of the nodes in *n*_*c*_ compared to the other nodes. In accordance with the formulation of NBS-SNI we propose in this paper, both groups of synthetic brain networks (condition and control) were induced the same high centrality nodes, i.e., the high centrality nodes in *n*_*c*_ were common to both groups of networks. The nodes in *n*_*c*_ serve as the important actors previously mentioned. Note that all connections are bidirectional, i.e., when a connection is added from node *u* to node *v*, the same connection is also added from node *v* to *u*. Note also, that an edge can only be added once, therefore the probabilities are adjusted accordingly during the generative process.

To obtain the parameters associated with *E*(*u, v*), we plotted the probability of connection *P*_*uv*_ between node *u* and *v* for the real structural network dataset of 264 brain regions (UCLA Autism; Rudie et al., 2013) as a function of the distance separating them *d*_*uv*_, considering both groups (ASD and control networks) altogether. We found that this relationship could be closely approximated by three different functions according to the range of distances. For distances ranging from *d*_*uv*_ = 0 up to *d*_*uv*_ = 18mm, a function a of the form 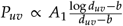 was fitted. The second part was approximated using a function of the form 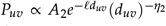 for distances ranging from 18 *≤ d*_*uv*_ < 39mm. For the last part (*d*_*uv*_ *≥* 39), a power law of the form *P*_*uv*_ ∝ *A*_3_(*d*_*uv*_)^*η* 3^ was fitted.

Since structural brain networks are generally represented as weighted networks, the distribution of weights from the real autism structural dataset was also extracted. We found that a function of the form *p*_*w*_ *≈ α*_*w*_ *· e*(^*kw*^)/*w* + *C*_*w*_, where *p*_*w*_ is the probability of finding a connection with weight *w* between two brain regions was a good approximation of the first half of this relationship. The second half (the right tail) of this relationship was approximated by a power law distribution 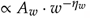, which allows for a few larger connection weights that cannot be accounted for by the first function (*A*_*w*_ *· e*(^*kw*^)/*w*).

For *E*(*u, v*), the parameters we found empirically are: (first part;*A*_1_ = 133.49 and *b* =*−*899.3), (second part; *A*_2_ = 0.000943, *ℓ* = 0.2277 and *η*_2_ = *−*3.83) and (third part; *A*_3_ =275.178 and *η*_3_ = 2.031). For the weights, we found *α*_*w*_ = 0.194, *k* = 1.038 *×* 10^*−*6^ and *C*_*w*_ = *−*0.0007 for the first part, and *A*_*w*_ = 627.78 and *η*_*w*_ = 2.99 for the second part. Moreover standard deviation 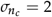 was kept constant for all our experiments.

### Functional network generative model

The synthetic functional brain networks are fully connected undirected weighted networks, i.e., a bidirectional weighted connection exists between every pair of nodes (*u, v*).

In the noise+contrast(s) group, the entries of the adjacency matrices can comprise different contrasts of connections on different sets of edges 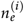 sampled from 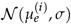, where (*i*) denotes the group in which the contrast(s) is present. The rest of the connection weights are sampled from *𝒩* (0, *σ*), thereby serving as noise. The probability density function for the weights thus is

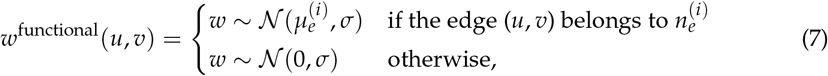

In the other group (noise only group) all the entries in the adjacency matrices are sampled from 𝒩 (0, *σ*). For *σ* = 1 (the value employed for all the functional contrasts in our simulations), this procedure has the effect of creating a contrast-to-noise ratio 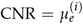 on the edges in 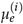 in one group, while the other group consists of adjacency matrices with all their entries sampled from Gaussian noise of zero mean and variance *σ*^2^.

### Numerical experiments

#### Identification of an embedded contrast

The synthetic networks for this experiment were built using the same nodes (*n* = 264; including their positions) as in the UCLA Autism dataset (Rudie et al., 2013), of which |*n*_*c*_| = 5 were chosen as being more central (see Fig. 4A). The same number of weighted edges, 3188, were randomly added using Eq. (5), with parameters 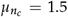 and 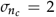. We generated 60 synthetic structural networks equally divided into the condition and the control groups, each network consisting in a single connected component.

**Figure 4.**
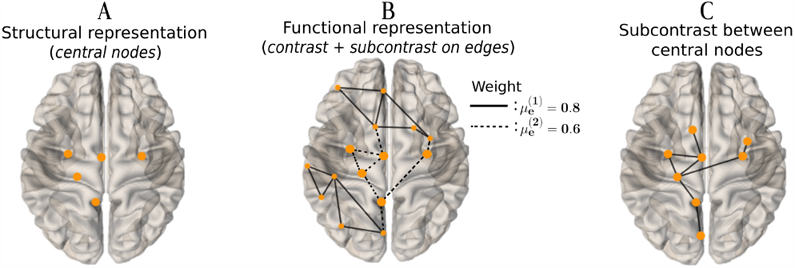
This synthetic data consists of network models of 264 nodes with 30 networks in each group. **A** Illustration of the 5 nodes in *n*_*c*_ on which a greater closeness centrality score was induced in the structural network models with 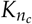(*u, v*) *∼ 𝒩* (1.5, 2). **B** Illustration of the functional connected component created around the 5 central nodes. Edges with a lower CNR sampled from *𝒩* (0.6, 1) are depicted as dashed lines, while edges belonging to the contrast of greater CNR sampled from *𝒩* (0.8, 1) are represented by solid lines. The five central nodes in *n*_*c*_ are depicted bigger. **C** The subcontrast, i.e., the connections of lower CNR interconnecting the 5 central nodes and reaching three other nodes placed at a maximum distance *d* = 1 from the central ones (now represented as solid lines). *Images generated with the brainconn R package* (https://github.com/sidchop/brainconn/)

To build the functional networks from the condition group, a first set of 8 edges connected to at least one node in *n*_*c*_ were given a weight using Eq. (7) with 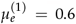 and *σ* = 1 (subcontrast edges; dashed lines in Fig. 4B). A second set of 11 edges located in the periphery of the 5 nodes in *n*_*c*_ were given a weight using Eq. (7) with 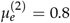and *σ* = 1 (contrast edges; solid lines in Fig. 4B). All other edges, as well as all edges in the functional networks belonging to the control group, were given a weight according to Eq. (7) (sampled from *𝒩* (0, 1)).

Our objective here is to illustrate how taking into account information about the nodes (here present in the structural networks) can allow the identification of edges that do not necessarily have the largest contrast-to-noise ratio (these subcontrast edges are shown in Figs. 4C and 5A). Figure 5 shows a comparison of the performances between NBS and NBS-SNI with *d* = 1.

**Figure 5.**
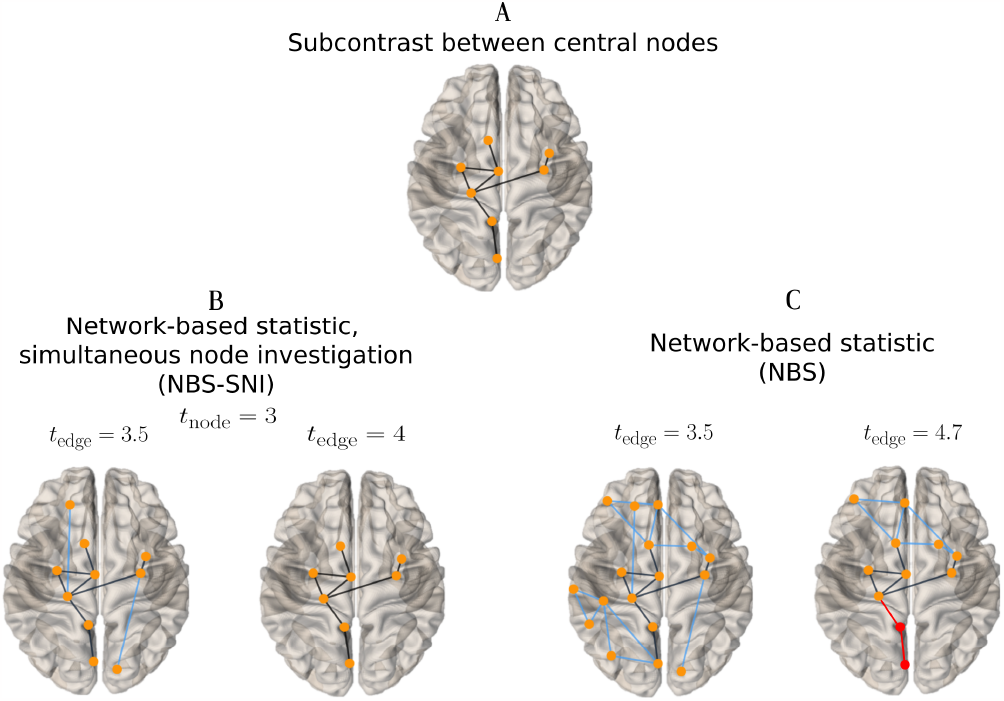
NBS-SNI and NBS performances at identifying a subcontrast were compared on the simulated data shown in Fig. 4, for two sets of parameter thresholds. Blue edges correspond to edges that were falsely identified by the method (false positives) and red edges correspond to edges that did belong to the contrast, but were not recovered by the method (false negatives). **A** The target subcontrast to be identified (a reproduction of Fig. 4C. **B** The subnetworks extracted with NBS-SNI across two different thresholds *t*_edge_. NBS-SNI perfectly recovers the simulated subcontrast. **C** The subnetwork extracted by NBS, which fails to perfectly capture the subcontrast, with 5 false positive and 2 false negative connections.

Blue edges correspond to connections identified by either method that did not belong to the subcontrast (false positives), while red edges correspond to connections that were missed (false negatives). Figure 5B shows the results for two sets of parameter thresholds *t*_edge_ obtained with NBS-SNI, which perfectly recovers the subcontrast edges. Figure 5C shows that *t*_edge_ could not be tuned for NBS to perfectly recover the subcontrast edges. For NBS, the threshold *t*_*edge*_ was increased to a larger value of 4.7 compared to NBS-SNI, where *t*_*edge*_ = 4. This is due to the fact that NBS-SNI is more restrictive than NBS, owing to the statistical test on the nodes. For NBS, this figure shows that past a threshold of 4.7, the method discards edges that belong to the contrast, even though it still misses some true positives.

Figure 5 illustrates what was already expected owing to the design of NBS-SNI, i.e., it can recover a subcontrast that is embedded inside another contrast of greater CNR, a task NBS is not suited for. Evidently, this conclusion is only valid if the nodes in the subcontrast share some common property (e.g., the closeness centrality in this example), which distinguishes them from the other nodes. In our example, this shared property between the nodes in the subcontrast was induced in the structural network models, but a nodal property could also be computed from the functional network representations. We chose this formulation since nodal properties based on shortest distances seem most suited for structural networks than for functional ones.

#### Systematic comparison

The next experiment aims at getting a better grasp of the cases when NBS-SNI offers a gain over NBS (and when it does not). Alongside NBS-SNI and NBS, we also consider the false discovery rate (FDR) to compare the two methods with a link-based controlling procedure.

Indeed, FDR treats every edge independently, as opposed to NBS and NBS-SNI, which cannot declare individual edges to be significant, but rather the component that they form. We evaluated the specificity and sensitivity of both NBS-SNI and NBS, along with the false discovery rate (FDR).

The experiment goes as follows. This procedure is repeated 1000 times for each set of parameters and thresholds (see Fig. 6 for details). Again, synthetic networks of 3188 edges were generated using the 264 nodes and their pairs of distances from the UCLA Autism dataset (Rudie et al., 2013). Among these nodes, |*n*_*c*_|= 5 are randomly selected to be the central nodes in the structural network. Among the 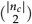 edges between these nodes and two other nodes (not present in *n*_*c*_) in the functional networks, a component of 20 edges consisting of a single contrast with mean CNR 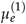 and variance *σ* = 1 is selected as the set 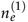. The component of functional edges therefore encompasses the five central nodes, plus two other nodes which do not possess higher induced centrality. Using Eqs. (5) and (7), 60 synthetic networks equally divided into the condition and the control groups are generated. (Recall that edges in the set 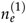 in the functional networks associated with the control group are no different than the other edges.) The most significant edges are then identified using the three methods (using *d* = 1 for NBS-SNI) and the true positive (TPR) and false positive rates (FPR) are computed. For NBS-SNI, *d* = 1 was used since some of the 20 connections in the contrast reach two nodes that do not belong to *n*_*c*_.

**Figure 6.**
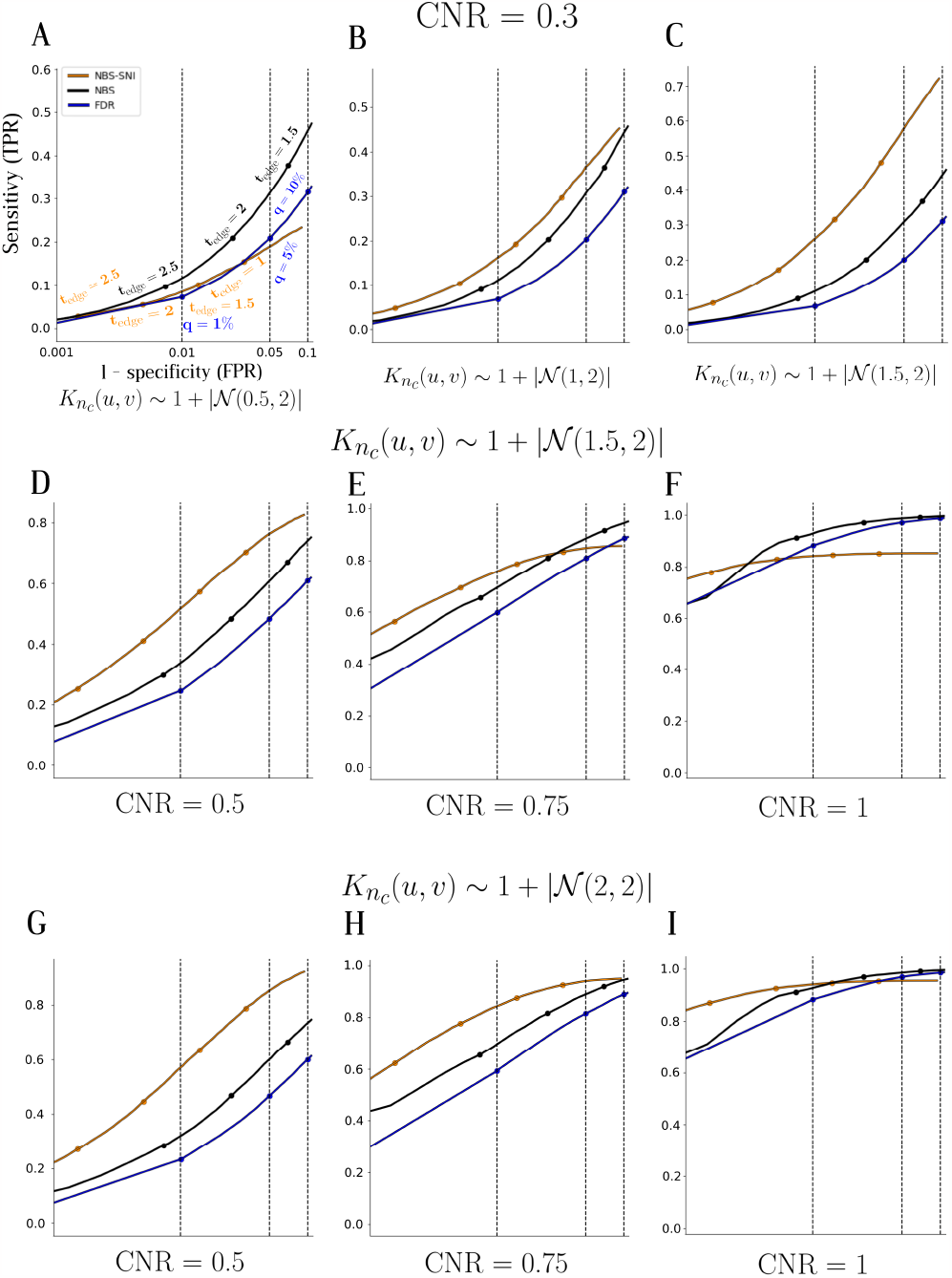
Receiver operating characteristic (ROC) curves were plotted to compare the sensitivity (**TPR**) and specificity (1 - **FPR**) of NBS-SNI (orange), NBS (black) and the false discovery rate (FDR) (blue), serving as a link-based controlling procedure, in detecting a simulated contrast on synthetic networks. Each point on a ROC curve for NBS-SNI and NBS (orange and black curves, respectively) represents the average of the TPR and FPR calculated from the 1000 realizations, at a specific parameter threshold *t*_edge_, some of which are indicated on the curves. For the FDR curves (in blue), each point represents a different *q*-value threshold, with a few indicated as a percentage. **A** ROC curves with the parameters 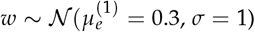 in the functional network models and 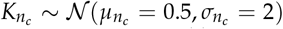 in the structural network models. **B** (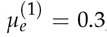; *σ* = 1) and 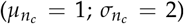. **C** (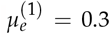; *σ* = 1) and (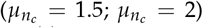). **D** 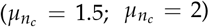 and (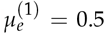; *σ* = 1) **E** 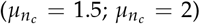 and (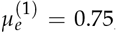; *σ* = 1). **F** 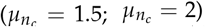 and (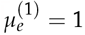; *σ* = 1). **G** 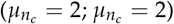 and (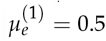; *σ* = 1). **H** 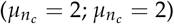 and (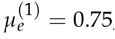; *σ* = 1). **I** 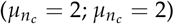 and (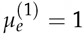; *σ* = 1).

The results are summarized by means of the receiver operating characteristic (ROC) curves shown in Fig. 6. In all cases, the value of the parameter thresholds ranged from *t*_edge_ = 0.5 to *t*_edge_ = 6 (for NBS-SNI and NBS) and *q* = 0.001 to *q* = 0.99 (for FDR; *q*-values represent the expected false discovery rate), with 100 equally spaced values in-between. For NBS-SNI, the parameter threshold for the nodes was kept constant at *t*_node_ = 0.5 for all realizations. For the synthetic functional networks, all the edge weights were sampled from a Gaussian distribution with with *μ* = 0 and *σ* = 1.

In Fig. 6A-C, the parameter 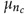 in 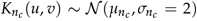 was varied in the structural network models for a given CNR = 0.3 (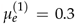; *σ* = 1) between the two groups in the functional network models. Since only the properties of the structural models are altered, NBS and FDR’s performances remain the same across Figs. 6A-C. It is only the performance of NBS-SNI that improves, i.e. the curves’ ascension is steeper, as the parameter 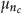 in the structural model is increased.

Figure 6D-F shows the ROC curves for three different CNRs in the functional network models, for a constant parameter 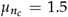 in the structural network models. The gain in statistical resolution offered by NBS-SNI compared to the others becomes less significant as the CNR is increased.

In Fig. 6G-I, the CNR was again varied, but for a constant structural model parameter increased to 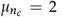. On the right, Fig. 6F with CNR = 1, all three methods offer similar performances after FPR = 0.01, thereby showing the gain in statistical resolution of NBSSNI and NBS over FDR to be most effective at lower contrast-to-noise ratios (Zalesky et al., 2010).

The following takeaway points from Fig. 6 pertain only to cases where the contrast forms a component and interconnects nodes that share a common property of interest (for e.g., high centrality nodes): (i) as the CNR (*μ*_*c*_) gets smaller, the better is the statistical resolution offered by NBS-SNI compared to NBS and FDR. (ii) The more pronounced is the node property of interest (parameter 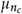) in the structural data, the greater is the statistical resolution offered by NBS-SNI. (iii) As the CNR is increased and 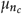 remains constant, the standard NBS and FDR will at some point offer a better resolution than NBS-SNI.

In sum, NBS-SNI offers a tighter control of the family-wise error rate than the other two methods in certain regimes by considering the nodes themselves, as opposed to considering only the edges. By giving importance to the nodes, the investigator can lower the parameter threshold *t*_edge_ to find significant abnormal connections of lower CNR. Conversely, the investigator could keep the same parameter threshold *t*_edge_ with NBS-SNI, thereby eliminating potential false positives that would result using NBS with the same value of *t*_edge_. This choice would depend on the desirable trade-off between finding true positives and eliminating false ones. It is also important to note that if the structural connectivity matrices, or the anatomical property of interest are noisy, or if a contrast of interest does not span across representations, NBS-SNI might lower the statistical resolution offered by the standard NBS.

In Fig. 7, we assessed the robustness of these conclusions by simulating structural connectivity matrices that might be more noisy, or of lower quality. We tested the case where NBS-SNI offered the greatest gain in statistical resolution in each row of Fig. 6, i.e., Fig. 6 C, D and G.

**Figure 7.**
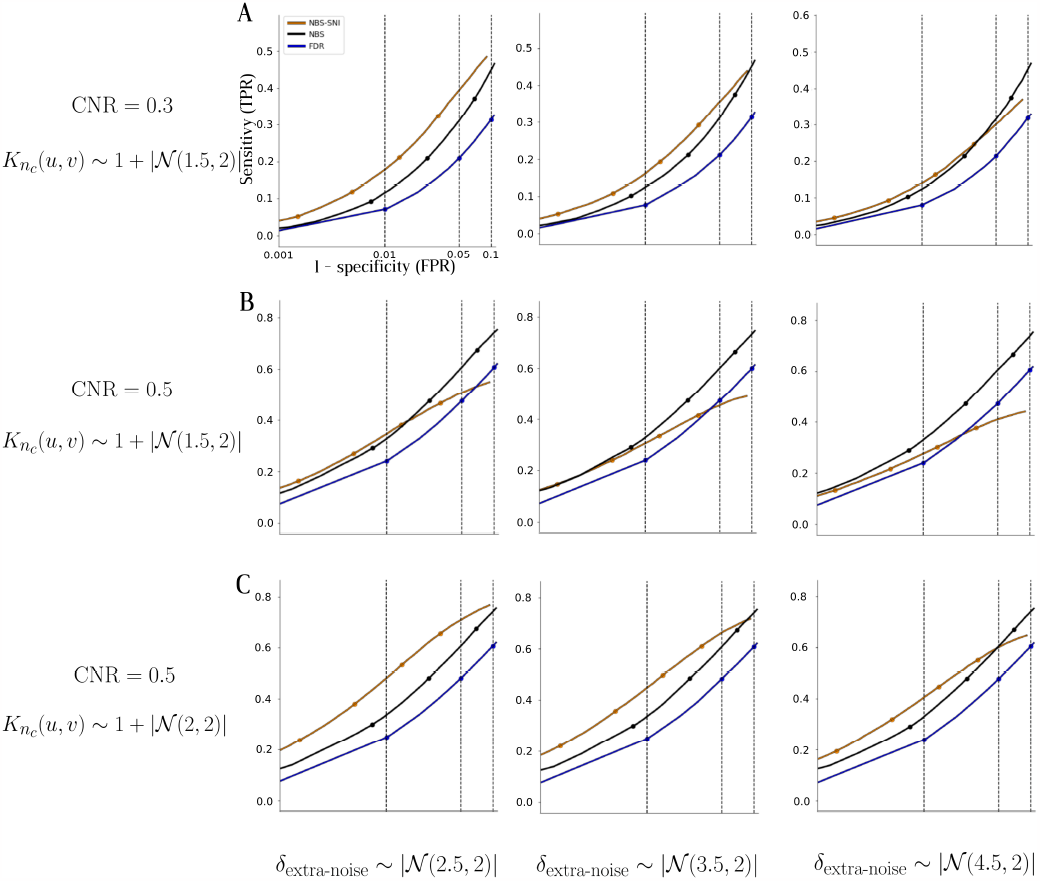
Receiver operating characteristic (ROC) curves were plotted to evaluate the robustness of the conclusions from Fig. 6, by adding noise to the probability distribution *P*^structural^(*u, v*) used to generate the synthetic networks. Each row shows the simulation results obtained for three different levels of extra noise, for a single pair of parameters (CNR; 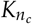 (*u, v*)). **A** Three different levels of noise added to the ROC curves of Fig. 6C. **B** Noise added to the case shown in Fig. 6D. **C** Noise added to the case shown in Fig. 6G.

For each pair of simulation parameters (CNR; 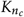 (*u, v*)), we added three different levels of noise *δ*_extra-noise_ to the probability distribution *P*^structural^(*u, v*) used to generate the structural connectivity matrices, i.e., *P* ^noisy-structural^(*u, v*) = *P*^structural^(*u, v*) + *δ*_extra-noise_. The approach was the following: we computed *P*^structural^(*u, v*) as before, but for each values *P*^structural^(*u, v*) we added noise terms sampled from three different folded normal distributions (i.e., taking the absolute values of the normal distribution to prevent including negative entries in the probability distribution). The three folded normal distributions we tested are *δ*_extra-noise_ *∼* |*𝒩* (2.5, 2)|, |*𝒩* (3.5, 2)| and |*𝒩* (4.5, 2)|.

The simulation results in Fig. 7 show that the gain in statistical resolution offered by NBS-SNI over NBS is most robust when the contrast-to-noise ratio (CNR) is low (Fig. 7A), or alternatively, when the contrast on the central nodes 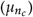 in the synthetic structural networks is high (Fig. 7C). In the middle row, the CNR is too high and 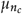 is too low, and the standard NBS performs better than NBS-SNI for most of the discrimination thresholds *t*_edge_. This example is a reminder for prospective user of the method that the extension we propose to the standard NBS is not intended as a replacement for the canonical method. Rather, we want to highlight the notion that in some regimes, NBS-SNI might offer a statistical gain over NBS. From these simulations, these regimes are characterized by a low CNR in the functional data, but also by an interplay of contrasts of interest in two different representations. In such cases, NBS-SNI can be used as a tool to leverage contrasts that span across representations.

The results presented in the ROC curves of Fig. 6 and 7 offer a somewhat narrow view of the discriminating performances of the different methods, owing to the specific parameters employed in each simulation. A more comprehensive interpretation of these results is given in Fig. 8, where the relative performances of NBS-SNI and NBS are shown when varying the number of samples and the overlap coefficient (OVL) between the Gaussian noise *𝒩* (0, 1) and the contrast 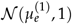 (the standard deviation was again kept constant at *σ* = 1 for all realizations). The values shown on the x-axis represents the number of synthetic samples in each group. The OVL measures the similarity between two distributions via the overlapping area of their distribution functions. As for the ROC curves in Fig. 6, *N* (*N*_1_ = *N*_2_) networks were generated in each group, with a contrast of 20 functional edges. Again, the position of the 264 nodes were taken from the from the UCLA Autism dataset (Rudie et al., 2013). Among these nodes, |*n*_*c*_| = 5 were randomly selected to be the central nodes in the structural network.

**Figure 8.**
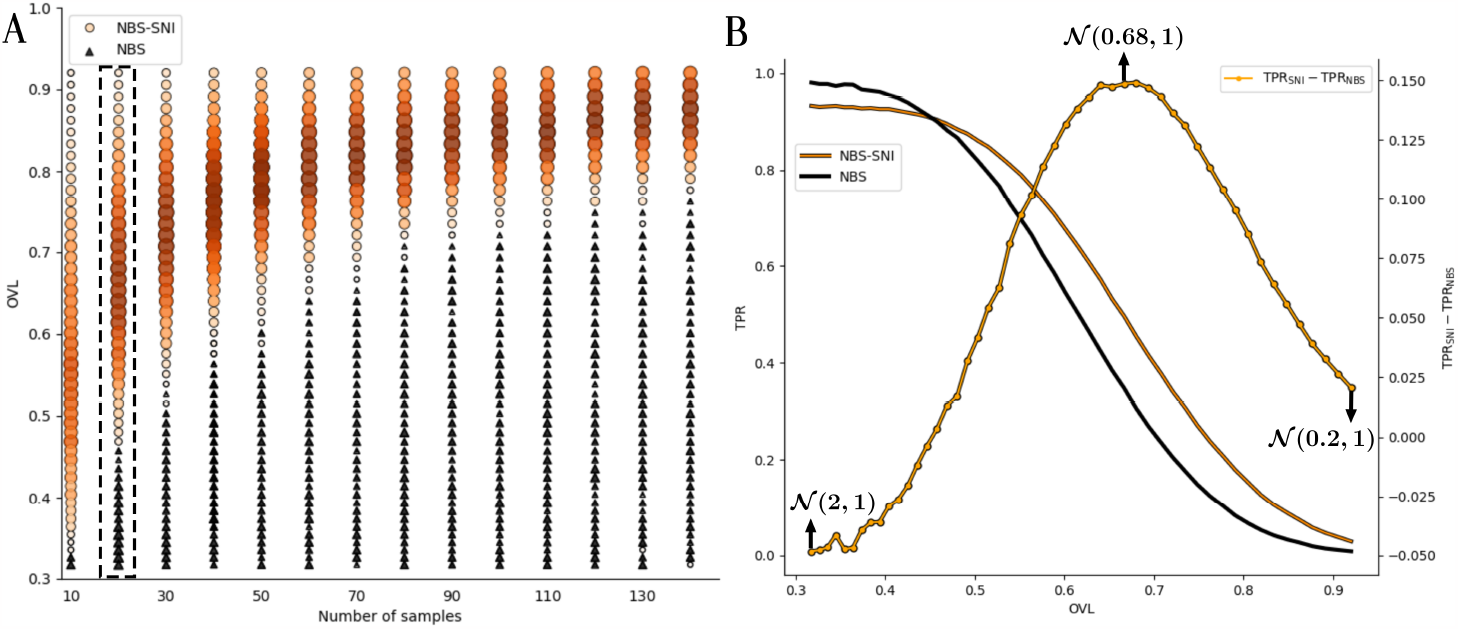
A Relative performance of NBS-SNI versus NBS at identifying a contrast of 20 edges in networks of 264 nodes as a function of the overlap coefficient (OVL) and the number of synthetic network samples generated in each group. Orange circles represent a set of parameters where NBS-SNI offered a greater true positive rate (TPR) than NBS, while black triangles show the cases where NBS yielded a greater TPR than NBS-SNI. For NBS-SNI, the size and color intensity of the points represents the relative gain in statistical resolution over NBS, calculated as TPR_SNI_*−* TPR_NBS_. A color map was not used for the cases where TPR_NBS_ *−*TPR_SNI_ for a clearer visualization; only the size of the black triangles reflects the gain in resolution. **B** TPR as a function of the overlap coefficient (OVL) for 20 samples, along with the curve showing the difference between the two performances, calculated as TPR_SNI_*−*TPR_NBS_. The orange (NBS-SNI) and black (NBS) curves correspond to the second column from the left in A.

To ensure a fair comparison of the true positive rate yielded by NBS-SNI and NBS, we controlled for the number of significant edges yielded by both methods. Their performances were compared based on subnetworks containing exactly 50 edges. To enforce this, the 200 (300) most significant t-scores with NBS (NBS-SNI) were retained and the largest connected component identified. For NBS, the top 50 most significant t-scores within the remaining connected component were retained. The same was done with NBS-SNI after taking the relaxed intersection (*d* = 1) with the central nodes (*t*_node_ = 0.5).

The orange circles represent a pair of parameters (number of samples, OVL) for which NBS-SNI offered a better performance (TPR_SNI_ *>* TPR_NBS_), where their sizes and colors reflect the relative gain in statistical resolution compared to NBS. The relative importance (in both color and size) of each point is calculated as TPR_SNI_ *−* TPR_NBS_. For visualization purposes, a colormap was not used for NBS, i.e. when TPR_NBS_ *>* TPR_SNI_ (black triangles); only the size of the triangles reflects the relative gain in statistical resolution offered by NBS over NBS-SNI. For every point (circle or triangle) in figure 8A, the simulation was repeated 500 times before the results were averaged to yield accurate visualization.

The results from Fig. 8 show that with kind of synthetic data, the range of CNR for which NBS-SNI offers a statistical resolution gain over NBS decreases with the number of samples, and that the gain is most important when the CNR is low (i.e., when the overlap coefficient is high). Altogether, the results figures 6 and 8 indicate that NBS-SNI can improve the statistical resolution in certain regimes by considering two representations simultaneously. More precisely, we expect NBS-SNI to outperform NBS at identifying a contrast of *relatively* small contrast-to-noise ratio (CNR) in the functional network representations, where the nodes comprised in this contrast manifest themselves as high centrality nodes in the structural network representations of both groups. The intuition for proposing this kind of synthetic data is that some brain regions (nodes) could manifest as important information carrier (central nodes) in the structural representation of both the condition and the control group, while the arrangement and the strength of the connections emanating from these central nodes may differ in the functional representation. In other words, the synthetic data model employed is such that the identity of some high centrality nodes is common to both groups (condition and control) in the structural networks, but the functional connections between theses nodes may vary across the two groups. By relatively small, we mean that the CNR is smaller than the standard deviation of the modelled Gaussian noise.

Such a configuration may allow us to be less restrictive on the threshold applied to the set of edges in the functional networks. Since NBS-SNI is also concerned with central nodes, which are retained from the statistical test applied to the structural network representations, acknowledging central nodes in our framework might help us throw out some false positive edges that could have been retained from the functional edges’ two-sample statistical test. We posit that the false positive rate can be lowered in such a case with NBS-SNI, while the true positive rate can be increased as seen in Fig. 8.

## 4 BRAIN NETWORK CLASSIFICATION PERFORMANCES OF NBS-SNI VS. NBS USING REAL CASE-CONTROL DATA

In this section, we present the results obtained with NBS-SNI applied to two real casecontrol studies. The first consists of individuals diagnosed with the autism spectrum disorder (ASD) and the second consists of individuals diagnosed with early psychosis (EP). All functional network representations are fully connected and weighted networks, where an edge represents the pairwise correlation between two brain regions. The procedure employed to yield predictions is detailed in section 2. All the machine learning algorithms paired with NBS-SNI and NBS used to perform predictions are the ones implemented by the Python package scikit-learn (Pedregosa et al., 2011), without any hyper-parameters optimization. In what follows, various machine learning (ML) algorithms were tested with the NBS-predict framework (Serin et al., 2021) for comparison purposes, but we want to emphasize that, depending on the research goal and the desired level of interpretability of the output networks, users should be mindful of which ML algorithms they base their findings on. Indeed, if interpretability of the outcome network is paramount to the research question, we recommend the investigators to utilize linear ML algorithms such as Linear Discriminant Analysis, Ridge Classifier, or Linear Support Vector Machine (SVM), as they allow for proper monitoring of the important edges (features) throughout the classification process.

### Autism dataset (ASD)

#### Sample characteristics

This dataset consists of 29 individuals diagnosed with ASD and 19 healthy individuals (see Table 1 for the demographics). Each individual in both groups is represented by a structural connectivity matrix and a functional connectivity matrix (resting-state). Moreover, there exists a node correspondence between all 264 nodes (mapped according to the Power264 brain atlas of Power et al. 2011) across both groups and both representations. These structural networks are sparse weighted and undirected networks, where an edge represents the density of white-mater axonal fiber tracts between two brain regions. All the data was collected using a Siemens 3T Trio scanner. For the fMRI data, resting-state data was collected during 6 minutes, with TR=3s, TE=28ms and 3x3x4 mm voxels. The structural data was obtained from 8 minutes DTI scans with 2x2x2mm voxels. This publicly available dataset was downloaded from the USC Multimodal Connectivity Database http://umcd.humanconnectomeproject.org.

**Table 1.** Demograhics of the UCLA autism dataset. Note that only the subjects for which both the functional and the structural connectivity matrices were available were retained from the larger dataset.

|  | Autism spectrum disorder<br>(ASD)<br>N = 29 | Typically developing<br>(TD)<br>N = 19 |
| --- | --- | --- |
| Baseline age, years ± SD | 13.2 ± 2.4 | 12.7 ± 1.9 |
| Baseline Females, N (%) | 4 (13.8) | 4 (21.1) |

## Results and Discussion

We used NBS-SNI with *d* = 1 (i.e., edges at a maximal distance of 1 from the central nodes are retained), paired with the machine learning and cross-validation procedure described earlier to perform classification of brain networks in a case-control setup. Brain networks are classified in either the ASD or control group. The parameters *K* = 15 and *r* = 25 were employed for the repeated k-fold cross-validation.

### Closeness centrality

The first node property investigated with NBS-SNI is the closeness centrality *C*_*C*_ (see section 2). The statistical test taking place on the measures of the nodes’ closeness centrality was the one-sample t-test. The classification results obtained with different machine learning algorithms are presented in Table 2. The best prediction performance was yielded by the Logistic Regression algorithm (*μ*_performance_ = 0.670 *±* 0.053) with the hyperconnected subnetwork, i.e., a hyperconnected subnetwork in the autism group of brain networks, containing an average of *k* _hypo_ = 188 *±* 20 functional connections across folds.

**Table 2.** Classification results obtained with NBS-SNI (*d* = 1) with the closeness centrality as the node property, with parameters *p*_node_ = 0.00001 and *t*_edge_ = 2. The average number of functional edges extracted by the method across folds are *k* _hyper_ = 188 *±* 20 and *k* _hypo_ = 121 *±* 19.

| Algorithms | | $\mu_{\text{performance}}$ | $\sigma_{\text{performance}}$ |
| --- | --- | --- | --- |
| Logistic Regression (L1) | hyper | **0.670** | **0.053** |
|  | hypo | 0.466 | 0.051 |
| K-Nearest Neighbors | hyper | 0.615 | 0.045 |
|  | hypo | 0.618 | 0.030 |
| Decision Tree | hyper | 0.600 | 0.072 |
|  | hypo | 0.478 | 0.062 |
| Ridge | hyper | 0.646 | 0.050 |
|  | hypo | 0.509 | 0.043 |
| SVM | hyper | 0.657 | 0.043 |
|  | hypo | 0.583 | 0.041 |
| Gaussian Naive Bayes | hyper | 0.623 | 0.049 |
|  | hypo | 0.596 | 0.035 |
| Stochastic Gradient Descent Classifier | hyper | 0.612 | 0.0400 |
|  | hypo | 0.540 | 0.053 |
| Linear Discriminant Analysis | hyper | 0.640 | 0.057 |
|  | hypo | 0.536 | 0.0570 |

For the hyperconnected case with the Logistic regression algorithm, applying a weight threshold of 1 to the final adjacency matrix containing the average prediction weight of every edges across folds as its entries, 23 hyperconnected edges remained. A hyperconnected component of 11 edges was identified within these 23 edges. The hyperconnected component is depicted in Fig. 9, alongside the list of its nodes and their associated degree. Interestingly, most of the connections are lateral (across hemispheres), with a larger proportion reaching the right putamen.

**Figure 9.**
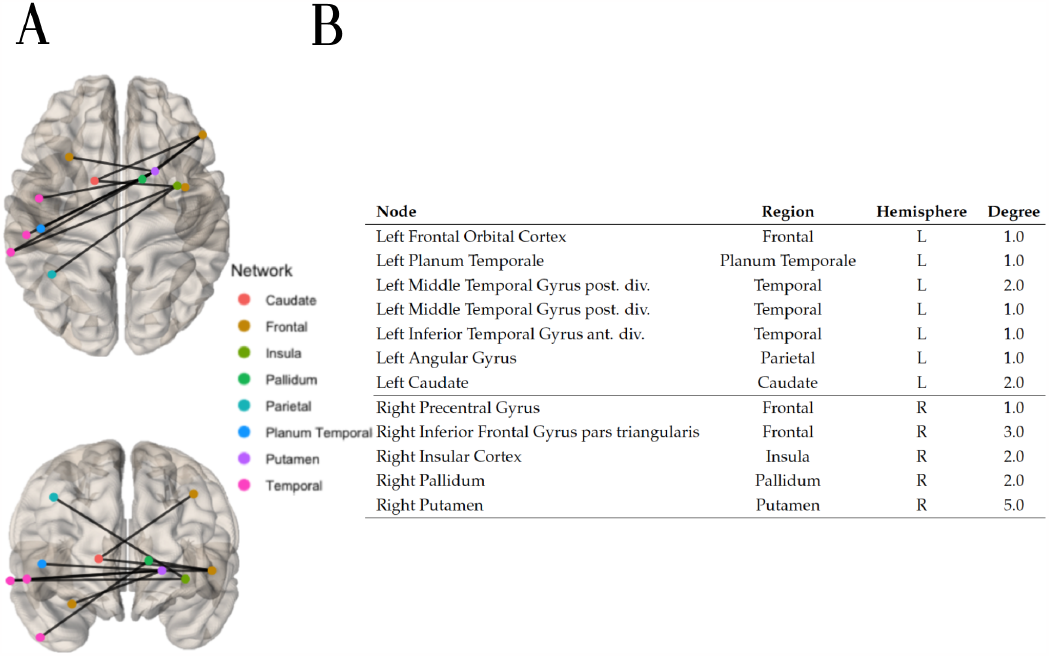
A Top and front views of the hypoconnected component extracted with the method NBS-SNI using the closeness centrality as the nodal property investigated and the Logistic regression algorithm to perform predictions. 23 edges which had a final prediction weight of 1 after scaling (0.673 before scaling) were retained. Within these 23 edges, the connected component of 11 edges shown was identified. **B** Nodal degree of the nodes present in the hypoconnected component on the left.

### Current flow closeness centrality

A variant of the closeness centrality, called the current flow closeness centrality (or sometimes information centrality) was also employed with NBS-SNI. As for the closeness centrality, the statistical test employed for this node property was the one-sample t-test. The prediction performances with this node property are presented in Table 3. The best performance was yielded by the Ridge classifier algorithm (*μ*_performance_ = 0.734 *±* 0.039) using the hypoconnected subnetwork, i.e., a disconnected subnetwork in the autism group of brain networks containing an average of *k* _hyper_ = 227 *±* 26 functional connections.

**Table 3.**
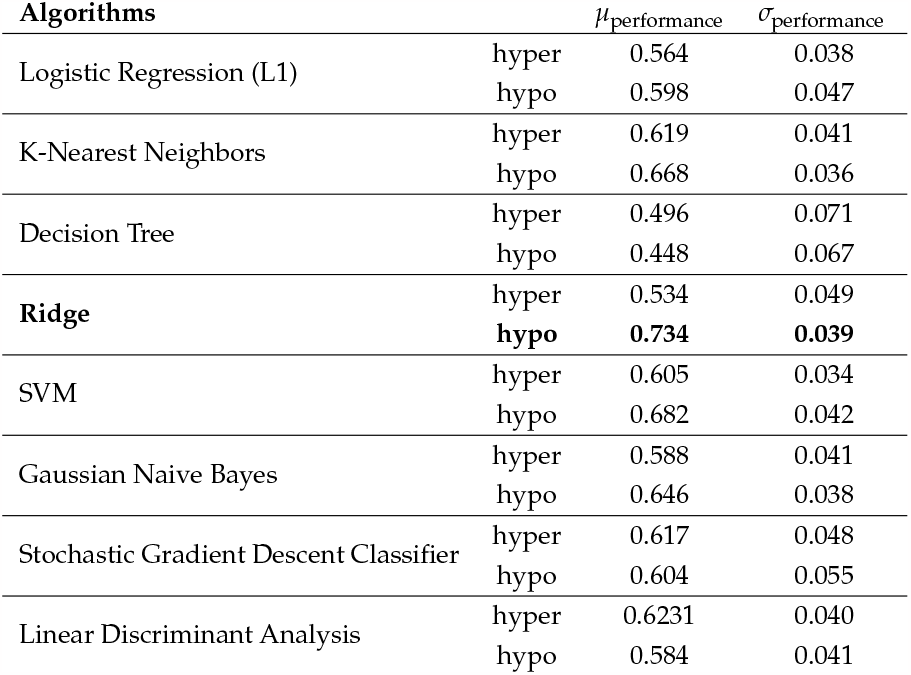
Classification results obtained with NBS-SNI (*d* = 1) with current flow closeness centrality as the node property, with parameters *p*_node_ = 0.000025 and *t*_edge_ = 2. The average number of functional edges extracted by the method across folds are *k* _hyper_ = 298 *±* 32 and *k* _hypo_ = 227 *±* 26.

| Algorithms | | $\mu_{\text{performance}}$ | $\sigma_{\text{performance}}$ |
| --- | --- | --- | --- |
| Logistic Regression (L1) | hyper | 0.564 | 0.038 |
|  | hypo | 0.598 | 0.047 |
| K-Nearest Neighbors | hyper | 0.619 | 0.041 |
|  | hypo | 0.668 | 0.036 |
| Decision Tree | hyper | 0.496 | 0.071 |
|  | hypo | 0.448 | 0.067 |
| **Ridge** | hyper | 0.534 | 0.049 |
|  | **hypo** | **0.734** | **0.039** |
| SVM | hyper | 0.605 | 0.034 |
|  | hypo | 0.682 | 0.042 |
| Gaussian Naive Bayes | hyper | 0.588 | 0.041 |
|  | hypo | 0.646 | 0.038 |
| Stochastic Gradient Descent Classifier | hyper | 0.617 | 0.048 |
|  | hypo | 0.604 | 0.055 |
| Linear Discriminant Analysis | hyper | 0.6231 | 0.040 |
|  | hypo | 0.584 | 0.041 |

Applying a weight threshold of 0.99 to the final prediction adjacency matrix (obtained via the Ridge classifier algorithm) yielded 43 hypoconnected edges. Within these 43 edges, a hypoconnected components of 23 edges was identified, which is shown in Fig. 10, alongside its nodes and their degree. Most connections seem to converge near the center of the sagittal plane of the brain. Notably is the large number of connections reaching the brainstem, the right cingulate gyrus posterior division and the right precuneous cortex. Together, these three brain regions account for 46% of all the connections implicated between all 20 brain regions of the hypoconnected component.

**Figure 10.**
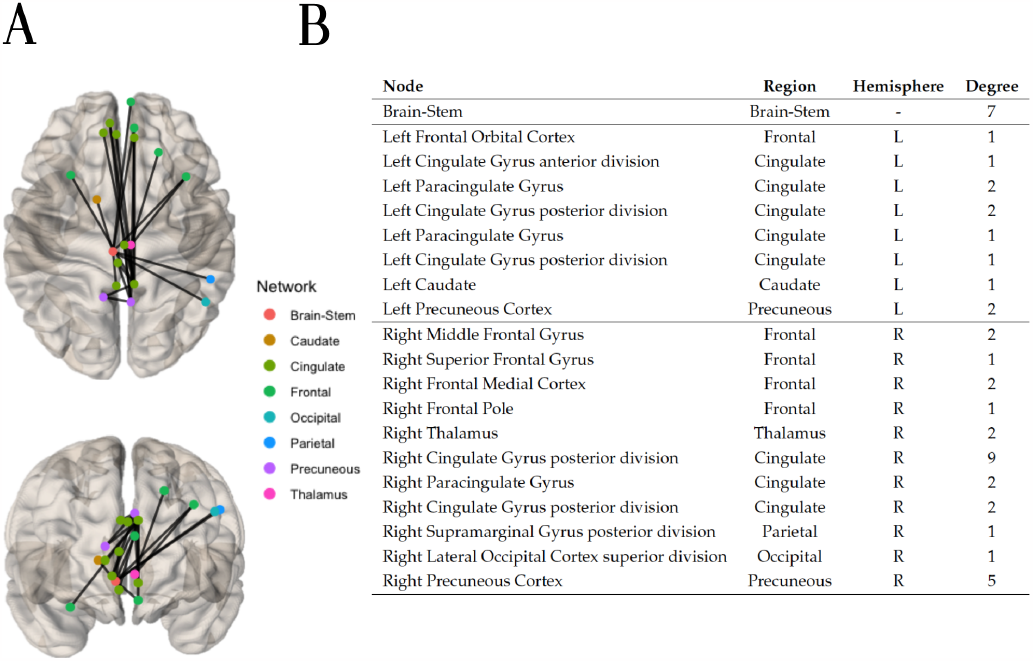
A Using the current flow closeness centrality, the hypoconnected subnetwork was extracted with the method NBS-SNI and the Ridge classifier algorithm. 43 edges which had a final prediction weight *≥*0.99 after scaling were retained. Within these 43 edges, the connected component of 23 edges shown (top and front views) was identified. **B** Nodal degree of the nodes present in the hypoconnected subnetwork on the left.

### Local information dimension (LID)

Finally the local information dimension was employed as the node property with NBSSNI. Due to the non-normal distribution of the computed nodes’ LID, a one-sample Kolmogorov–Smirnov test (K-S test) was used for this node property. To employ this node measure with the weighted networks of the structural representations, the sizes of the box *l* around a node was discretized into 15 equally spaced parts between the minimum distance and half of the maximum weighted distance between two nodes within the network. The prediction performances are presented in Table 4. The best performance was yielded by Decision tree classifier algorithm (*μ*_performance_ = 0.704 *±* 0.063), using the hyperconnected subnetwork, containing on average *k* _hyper_ = 124 *±* 18 functional connections.

**Table 4.** Classification results obtained with NBS-SNI (*d* = 1) with local information dimension (LID) as the node property, with parameters *p*_node_ = 0.005 and *t*_edge_ = 2.15. The average number of functional edges extracted by the method across folds are *k* _hyper_ = 124 *±* 18 and *k* _hypo_ = 98 *±* 30.

| Algorithms | | $\mu_{\text{performance}}$ | $\sigma_{\text{performance}}$ |
| --- | --- | --- | --- |
| Logistic Regression (L1) | hyper | 0.545 | 0.058 |
|  | hypo | 0.565 | 0.0451 |
| K-Nearest Neighbors | hyper | 0.592 | 0.058 |
|  | hypo | 0.623 | 0.051 |
| **Decision Tree** | **hyper** | **0.704** | **0.063** |
|  | hypo | 0.617 | 0.08 |
| Ridge | hyper | 0.629 | 0.036 |
|  | hypo | 0.501 | 0.07 |
| SVM | hyper | 0.604 | 0.044 |
|  | hypo | 0.599 | 0.053 |
| Gaussian Naive Bayes | hyper | 0.603 | 0.040 |
|  | hypo | 0.603 | 0.050 |
| Stochastic Gradient Descent Classifier | hyper | 0.566 | 0.041 |
|  | hypo | 0.554 | 0.053 |
| Linear Discriminant Analysis | hyper | 0.610 | 0.051 |
|  | hypo | 0.545 | 0.055 |

Applying a weight threshold of 0.99 to the final prediction adjacency matrix yielded by the Decision tree classifier algorithm resulted in 17 hyperconnected edges. Within these 17 edges, the hyperconnected component of 10 edges was identified and is depicted in Fig. 11, alongside the nodes and their degree. The majority of the nodes involved in the hyperconnected subnetwork are localized in the right hemisphere, connecting regions located solely in the frontal, temporal and occipital parts of the brain.

**Figure 11.**
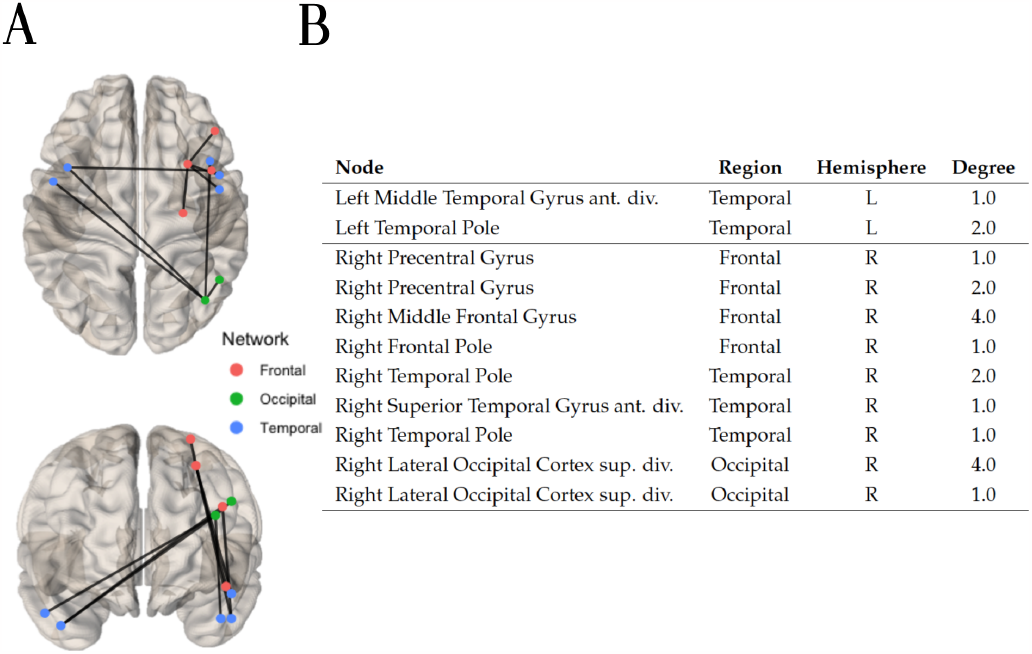
A Using the local information dimension (LID) the hyperconnected subnetwork was extracted with the method NBS-SNI and the Decision tree classifier algorithm. 17 edges which had a final prediction weight *≥*0.99 after scaling were retained. Within these 17 edges, the connected component of 10 edges was identified. **B** Nodal degree in the hyperconnected subnetwork depicted on the left.

### Comparison of predictions performances with NBS

Finally, the standard NBS was also employed to perform predictions. A range of parameter thresholds *t*_edge_ was tried, with *t*_edge_ = 2.5 being the one that yielded the best prediction performance. The classification results are presented in Table 5. The best prediction performance was yielded by the Gaussian Naive Bayes algorithm *μ*_performance_ = 0.639 *±* 0.026 with the hypoconnected subnetwork, containing on average *k* _hypo_ = 319 *±* 33 edges.

**Table 5.** Classification results obtained with the standard NBS with parameter *t*_edge_ = 2.5. The average number of functional edges extracted by the method across folds are *k* _hyper_ = 347 *±* 37 and *k* _hypo_ = 319 *±* 33.

| Algorithms | | $\mu_{\text{performance}}$ | $\sigma_{\text{performance}}$ |
| --- | --- | --- | --- |
| Logistic Regression (L1) | hyper | 0.461 | 0.033 |
|  | hypo | 0.570 | 0.026 |
| K-Nearest Neighbors | hyper | 0.606 | 0.039 |
|  | hypo | 0.630 | 0.031 |
| Decision Tree | hyper | 0.547 | 0.063 |
|  | hypo | 0.510 | 0.059 |
| Ridge | hyper | 0.554 | 0.052 |
|  | hypo | 0.562 | 0.044 |
| SVM | hyper | 0.598 | 0.033 |
|  | hypo | 0.615 | 0.031 |
| **Gaussian Naive Bayes** | hyper | 0.604 | 0.042 |
|  | **hypo** | **0.639** | **0.026** |
| Stochastic Gradient Descent Classifier | hyper | 0.5364 | 0.066 |
|  | hypo | 0.530 | 0.043 |
| Linear Discriminant Analysis | hyper | 0.552 | 0.053 |
|  | hypo | 0.586 | 0.042 |

To assess the significance of the improved prediction performance of NBS-SNI over NBS, permutations of individuals’ group memberships were performed. The prediction performances of both methods were calculated for 1000 permutations. For each permutation, the group membership of every individual was randomly assigned before performing the nested cross validation procedure. The p-values were calculated as the number of times the maximum accuracy with random group memberships 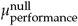 was greater or equal to the empirical accuracy *μ*_performance_, divided by the number of permutations (1000). The results presented in Fig. 12 show that the prediction performance of 74% yielded by NBSSNI using the current flow closeness centrality was found to be statistically significant (*p* = 0.016) under the permutation test. For NBS, the accuracy of 64% was not deemed significant (*p* = 0.122).

**Figure 12.**
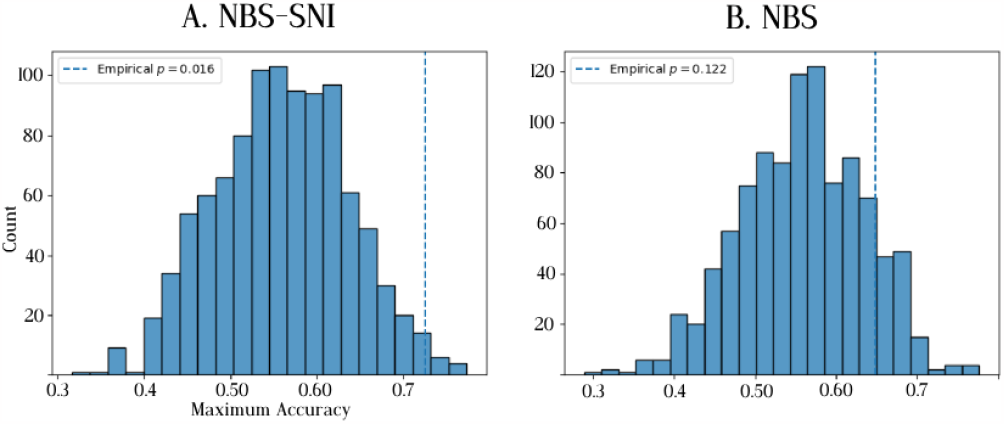
Distribution of maximum accuracy when subjects are randomly assigned to a group. A total of 1000 permutations were performed. The vertical dashed line represents the empirical prediction accuracy *μ*_performance_ obtained with the appropriate group memberships. The p-values are calculated as 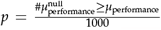. **A** The prediction accuracy of 74% obtained with NBS-SNI, using the current flow closeness centrality of nodes was deemed significant according to this permutation test. **B** The prediction accuracy of 64% obtained with NBS was not deemed significant.

The null distribution of prediction performances under group permutations is not centered at 50%, as one would expect. This indicates that prediction performances below *∼* 70% should be taken with a grain of salt with this dataset. This could be due to an artificial bias resulting from the small sample size of this dataset. Furthermore, this effect (i.e., a non-negligible proportion of null prediction performances greater than 50%) is most pronounced with NBS in Fig. 12B, therefore potentially showing a greater statistical discrimination offered by NBS-SNI with this dataset.

### HCP-Early Psychosis (EP)

#### Sample characteristics

This dataset consists of 75 individuals diagnosed with non-affective psychosis and 45 healthy control individuals (see Table 6 for the demographics) obtained from the Human Connectome Project (D. C. Van Essen et al., 2013). The fMRI data was parcellated using the Schaefer 200 7 networks parcellation (Schaefer et al., 2018). The anatomical measures of intrinsic curvature index and gray matter volume employed with NBS-SNI were calculated with FreeSurfer. The protocol scan sequences are: T1w (MPRAGE) and T2w (SPACE) structural scans of 0.8mm isotropic resolution. For the resting-state functional data, scans were obtained at 2mm isotropic resolution, multiband (MB) acceleration factor of 8 and TR 720ms, acquired twice: once with AP and once with PA phase encoding.

**Table 6.**
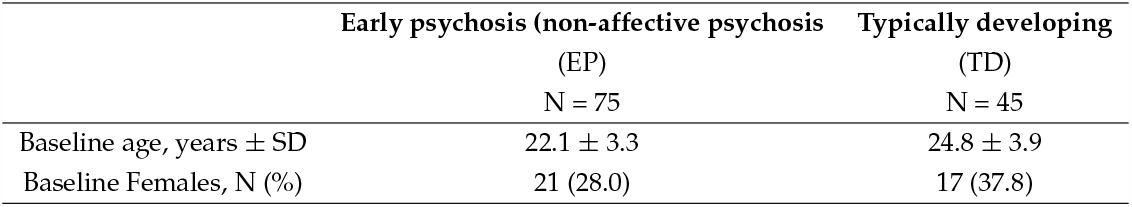
Demograhics of the HCP early psychosis dataset. Only the patients presenting a diagnosis of non-affective psychosis (N = 75) were retained from the original patient dataset (N= 88 EP).

|  | Early psychosis (non-affective psychosis<br>(EP)<br>N = 75 | Typically developing<br>(TD)<br>N = 45 |
| --- | --- | --- |
| Baseline age, years ± SD | 22.1 ± 3.3 | 24.8 ± 3.9 |
| Baseline Females, N (%) | 21 (28.0) | 17 (37.8) |

## Results and Discussion

For this dataset, the anatomically derived measures of intrinsic curvature index and gray matter volume were employed as the node property within the NBS-SNI framework. In this case, the statistical test can be performed directly on the anatomical measure to probe for abnormal functional connections between certain anatomical regions. The parameters *K* = 15 and *r* = 10 were employed for the repeated k-fold cross-validation in the following results.

### NBS-SNI with anatomical node property: intrinsic curvature index (ICI)

For the intrinsic curvature index (ICI), two independent one sample t-tests were performed with the parameter *p*_node_ = 0.20, and the intersection between the two sets of nodes was retained. For the hypoconnected edges, the two sample t-test parameter threshold was set to *t*_edge_ = 4.2. For the hyperconnected edges, this parameter had to be set to its minimal value *t*_edge_ = 0 in order to retain at least a few edges across all folds during the cross-validation process. This indicates that the early psychosis group had very few hyperconnected edges, while the control group had many more stronger functional connections. Once again, the parameter *d* = 1 was employed with NBS-SNI when taking the intersection between the nodes and the functional edges. The prediction performances are presented in Table 7. The best performance was yielded by Ridge classifier algorithm (*μ*_performance_ = 0.762 *±* 0.015), using the hypoconnected subnetwork, containing on average *k* _hypo_ = 620 *±* 200 functional connections.

**Table 7.** Classification results obtained with NBS-SNI (*d* = 1) using the anatomical measure of intrinsic curvature index as the node property, with parameters *p*_node_ = 0.20. For this dataset, the parameter threshold on the edges for the hyperconnected subnetwork had to be set to *t*_edge_ = 0 in order to retrieve at least a few hyperconnected edges after the statistical tests, while the optimal parameter for the hypoconnected subnetwork was found to be *t*_edge_ = 4.2. The average number of functional edges extracted by the method across folds are *k* _hyper_ = 292 *±* 75 and *k* _hypo_ = 620 *±* 200.

| Algorithms | | $\mu_{\text{performance}}$ | $\sigma_{\text{performance}}$ |
| --- | --- | --- | --- |
| Logistic Regression (L1) | hyper | 0.623 | 0.004 |
|  | hypo | 0.702 | 0.011 |
| K-Nearest Neighbors | hyper | 0.603 | 0.022 |
|  | hypo | 0.692 | 0.019 |
| Decision Tree | hyper | 0.533 | 0.047 |
|  | hypo | 0.665 | 0.040 |
| **Ridge** | hyper | 0.588 | 0.033 |
|  | **hypo** | **0.762** | **0.015** |
| SVM | hyper | 0.625 | 0.001 |
|  | hypo | 0.695 | 0.013 |
| Gaussian Naive Bayes | hyper | 0.550 | 0.015 |
|  | hypo | 0.685 | 0.011 |
| Stochastic Gradient Descent Classifier | hyper | 0.543 | 0.034 |
|  | hypo | 0.749 | 0.026 |
| Linear Discriminant Analysis | hyper | 0.543 | 0.038 |
|  | hypo | 0.715 | 0.015 |

After applying a weight threshold of 1, due to the large number of edges still remaining, only the top 10 nodes with the greatest degree were retained (corresponding to the 95^th^ percentile) and the hypoconnected component of 16 edges depicted in Fig. 13 was identified.

**Figure 13.**
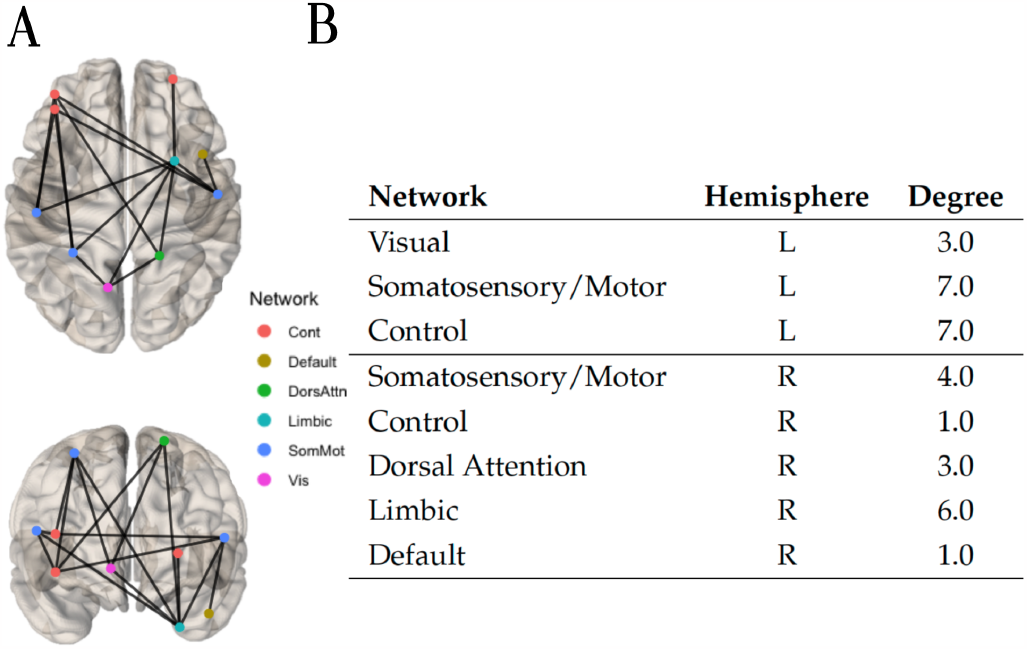
A Using the anatomical property of curvature index, the hypoconnected subnetwork was extracted with the method NBS-SNI and the Ridge classifier algorithm. Edges with a final prediction weight of 0.762 were retained. Due to the large number of edges possessing a final prediction weight of 1 (after scaling), only the top 10 nodes with the greatest degree (corresponding to the 95^th^ percentile) were retained. All the remaining edges already formed the connected component of 16 edges (top and front view) shown. **B** Nodal degree and network membership within the 7 networks Schaefer 200 parcellation.

### NBS-SNI with anatomical node property: gray matter volume

Using the gray matter volume (GMV) as a node property, two independent one sample ttests were performed with the parameter *p*_node_ = 0.001, and the intersection between the two sets of nodes was retained. For the hypoconnected edges, the two sample t-test parameter threshold was set to *t*_edge_ = 4.2. Once again, the parameter *d* = 1 was employed with NBS-SNI when taking the intersection between the nodes and the functional edges. The prediction performances are presented in Table 8. The best performance was yielded by Ridge classifier algorithm (*μ*_performance_ = 0.774 *±* 0.029), using the hypoconnected subnetwork, containing on average *k* _hypo_ = 640 *±* 191 functional connections.

**Table 8.** Classification results obtained with NBS-SNI (*d* = 1) using the anatomical measure of gray matter volume as the node property, with parameters *p*_node_ = 0.001. For this dataset the parameter threshold on the edges for the hyperconnected subnetwork had to be set to *t*_edge_ = 0 in order to retrieve at least a few hyperconnected edges after the statistical tests, while the optimal parameter for the hypoconnected subnetwork was found to be *t*_edge_ = 4.2. The average number of functional edges extracted by the method across folds are *k* _hyper_ = 273 *±* 60 and *k* _hypo_ = 640 *±* 191.

| Algorithms | | $\mu_{\text{performance}}$ | $\sigma_{\text{performance}}$ |
| --- | --- | --- | --- |
| Logistic Regression (L1) | hyper | 0.625 | 0.004 |
|  | hypo | 0.700 | 0.007 |
| K-Nearest Neighbors | hyper | 0.592 | 0.026 |
|  | hypo | 0.678 | 0.016 |
| Decision Tree | hyper | 0.533 | 0.047 |
|  | hypo | 0.693 | 0.045 |
| **Ridge** | hyper | 0.548 | 0.044 |
|  | **hypo** | **0.774** | **0.029** |
| SVM | hyper | 0.625 | 0.001 |
|  | hypo | 0.691 | 0.013 |
| Gaussian Naive Bayes | hyper | 0.543 | 0.013 |
|  | hypo | 0.683 | 0.010 |
| Stochastic Gradient Descent Classifier | hyper | 0.538 | 0.05 |
|  | hypo | 0.762 | 0.022 |
| Linear Discriminant Analysis | hyper | 0.565 | 0.028 |
|  | hypo | 0.713 | 0.020 |

After applying a weight threshold of 1, due to the large number of edges still remaining, only the top 11 nodes with the greatest degree were retained (corresponding to the 95_th_ percentile) and the hypoconnected component of 20 edges depicted in Fig. 14 was identified.

**Figure 14.**
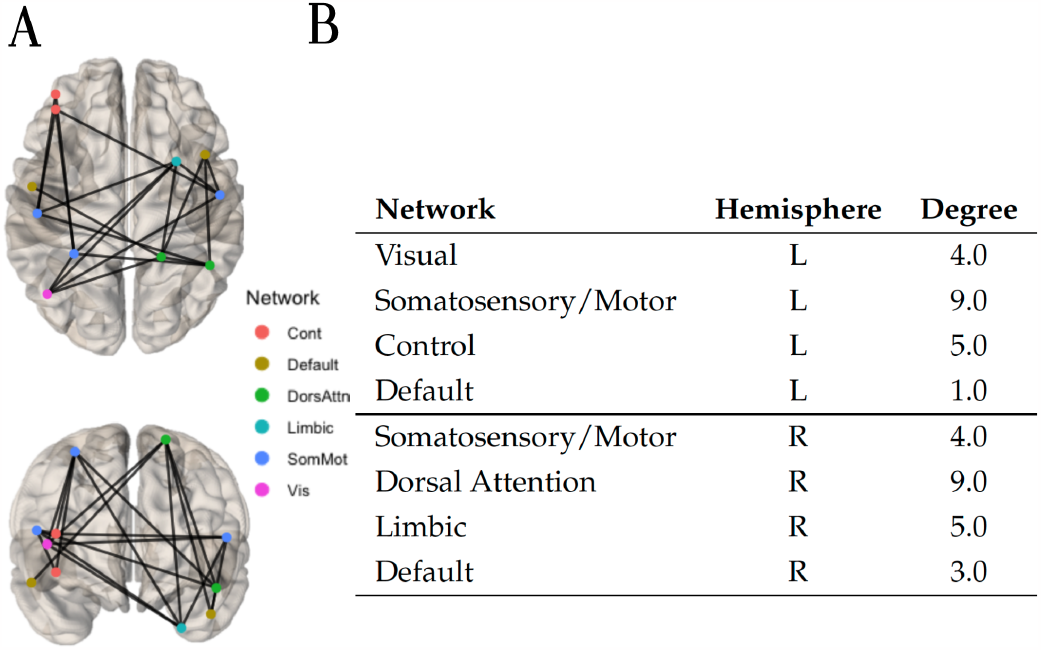
A Using the anatomical property of gray matter volume, the hypoconnected subnetwork was extracted with the method NBS-SNI (*d* = 1) and the Ridge classifier algorithm. Edges with a final prediction weight of 0.774 were retained. Due to the large number of edges possessing a final prediction weight of 1 (after scaling), only the top 11 nodes with the greatest degree (corresponding to the 95^th^ percentile) were retained. All the remaining edges already formed the connected component of 20 edges (top and front view) shown. **B** Nodal degree and network membership within the 7 networks Schaefer 200 parcellation.

In the following results, a two-sample t-test was applied to the node property of gray matter volume, rather than two independent one-sample t-tests, as it was done for the previous results. Applying a two-sample t-test on the node properties implies in this case that NBS-SNI probes for differences in gray matter volume across the condition and the control group, rather than searching for nodes that have high values of gray matter volume in both groups (i.e., resulting from taking the intersection of the suprathreshold nodes from both one-sample t-tests). The reason for this choice was motivated by the large body of literature reporting loss of gray matter volume in individuals with schizophrenia (Gur et al., 2000, 1999; Hulshoff Pol et al., 2002; Vita et al., 2012). Applying a two-sample t-tests on the node to investigate gray matter volume abnormalities between groups therefore seems appropriate.

NBS-SNI was employed with the parameters 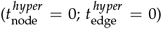 for the hyperconnected case and 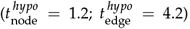. The prediction performances are presented in Table 9. The best performance was yielded by Ridge classifier algorithm (*μ*_performance_ = 0.776 *±* 0.021), using the hypoconnected subnetwork, containing on average *k* _hypo_ = 903 *±* 301 functional connections.

**Table 9.**
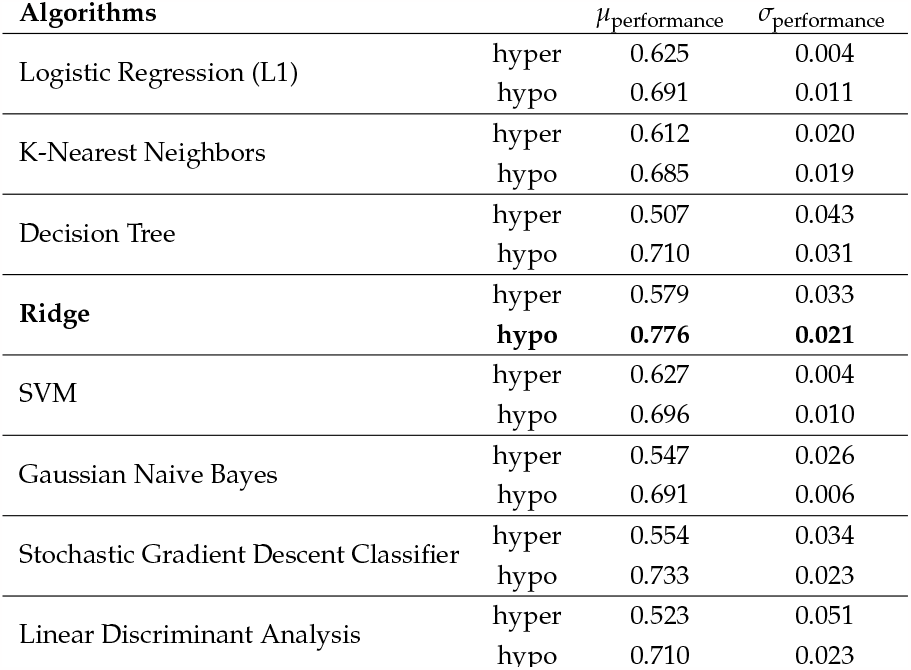
Classification results obtained with NBS-SNI (*d* = 1) using the anatomical measure of gray matter volume as the node property, with parameter thresholds 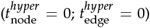 for the hyperconnected case and 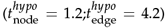 for the hypoconnected one. For this investigation, a two-sample t-test was applied to the node property of gray matter volume. The average number of functional edges extracted by the method across folds are *k* _hyper_ = 119 *±* 52 and *k* _hypo_ = 903 *±* 301.

After applying a weight threshold of 1, due to the large number of edges still remaining, only the top 13 nodes with the greatest degree were retained (corresponding to the 95^th^ percentile) and the hypoconnected component of 22 edges depicted in Fig. 15 was identified. This hypoconnected subnetwork shows connections that were weaker (hypo), but also between nodes of lower gray matter volume in the early psychosis group. Using our framework, a thorough investigation of this condition could therefore investigate how functional connectivity is related with gray matter volume.

**Figure 15.**
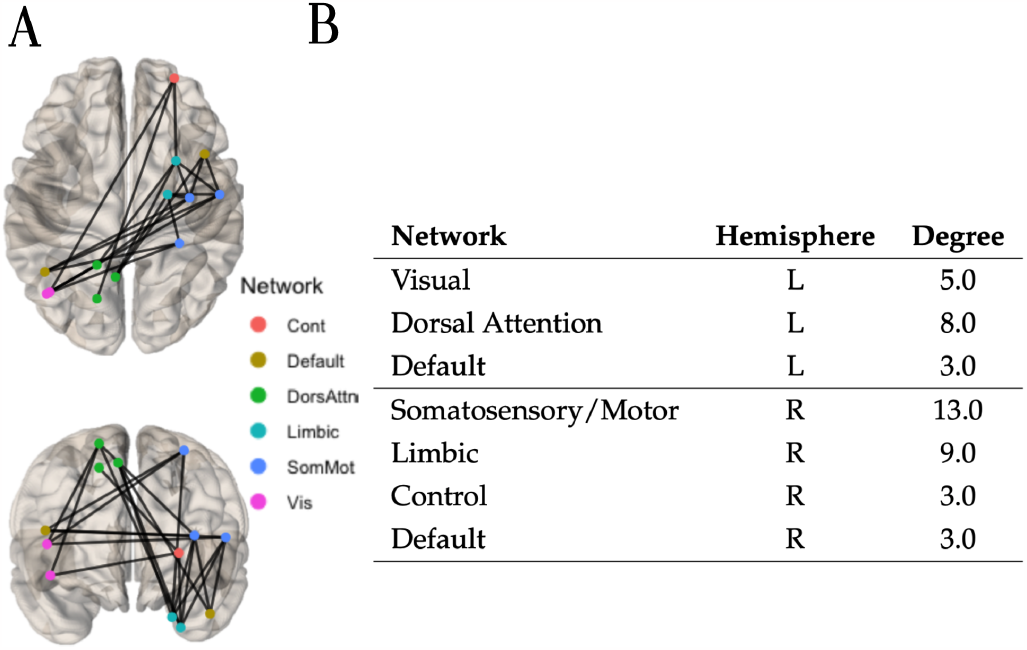
A Using the anatomical property of gray matter volume and a two-sample t-test on the node to probe for anatomical regions of reduced gray matter volume, the hypoconnected subnetwork was extracted with the method NBS-SNI and the Ridge classifier algorithm. Edges with a final prediction weight of 0.776 were retained. Due to the large number of edges possessing a final prediction weight of 1 (after scaling), only the top 13 nodes with the greatest degree (corresponding to the 95^th^ percentile) were retained. All the remaining edges already formed the connected component of 22 edges (top and front view) shown. **B** Nodal degree and network membership within the 7 networks Schaefer 200 parcellation.

### Comparison of predictions performances with NBS

Finally, the standard NBS was also employed to perform predictions. A range of parameter thresholds *t*_edge_ was tried, with *t*_edge_ = 2.5 being the one that yielded the best prediction performance. The classification results are presented in Table 10. The best prediction performance was yielded by the Ridge classifier algorithm *μ*_performance_ = 0.727 *±* 0.013 with the hypoconnected subnetwork, containing on average *k* _hypo_ = 9330 *±* 990 edges.

**Table 10.** Classification results obtained with the standard NBS with parameter *t*_edge_ = 0 for the hyperconnected case and *t*_edge_ = 2.5 for the hypoconnected one. The average number of functional edges extracted by the method across folds are *k* _hyper_ = 436 *±* 116 and *k* _hypo_ = 9330 *±* 990.

| Algorithms | | $\mu_{\text{performance}}$ | $\sigma_{\text{performance}}$ |
| --- | --- | --- | --- |
| Logistic Regression (L1) | hyper | 0.619 | 0.006 |
|  | hypo | 0.672 | 0.01 |
| K-Nearest Neighbors | hyper | 0.593 | 0.02 |
|  | hypo | 0.664 | 0.015 |
| Decision Tree | hyper | 0.543 | 0.030 |
|  | hypo | 0.668 | 0.035 |
| **Ridge** | hyper | 0.530 | 0.026 |
|  | **hypo** | **0.727** | **0.013** |
| SVM | hyper | 0.625 | 0.001 |
|  | hypo | 0.683 | 0.011 |
| Gaussian Naive Bayes | hyper | 0.523 | 0.035 |
|  | hypo | 0.674 | 0.005 |
| Stochastic Gradient Descent Classifier | hyper | 0.512 | 0.032 |
|  | hypo | 0.722 | 0.032 |
| Linear Discriminant Analysis | hyper | 0.572 | 0.033 |
|  | hypo | 0.703 | 0.016 |

After applying a weight threshold of 1, due to the large number of edges still remaining, only the top 10 nodes with the greatest degree were retained (corresponding to the 95^th^ percentile) and the hypoconnected component of 42 edges depicted in Fig. 16 was identified.

**Figure 16.**
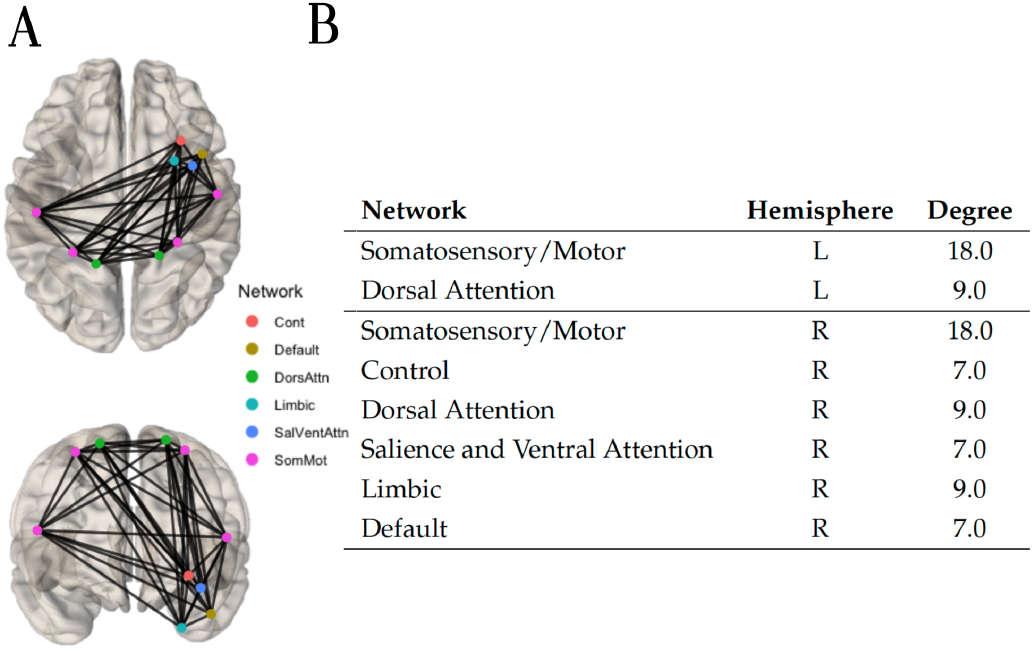
A The hypoconnected subnetwork was extracted with the standard NBS and the Ridge classifier algorithm. Edges with a final prediction weight of 0.727 were retained. Due to the large number of edges possessing a final prediction weight of 1 (after scaling), only the top 10 nodes with the greatest degree were retained (corresponding to the 95^th^ percentile). All the remaining edges already formed the connected component of 42 edges (top and front view) shown. **B** Nodal degree and network membership within the 7 networks Schaefer 200 parcellation.

The results from the permutation test are presented in Fig. 17. Overall, the prediction accuracies yielded by NBS-SNI with different anatomical properties and the standard NBS were all deemed significant by this permutation test. When employed with gray matter volume (GMV) as a node property, using a two-sample t-test on the nodes’ statistical test, NBS-SNI offered a marginal gain of five percentage points of the standard NBS. In all cases, there was no overlap between the empirical prediction accuracy and its null distribution.

**Figure 17.**
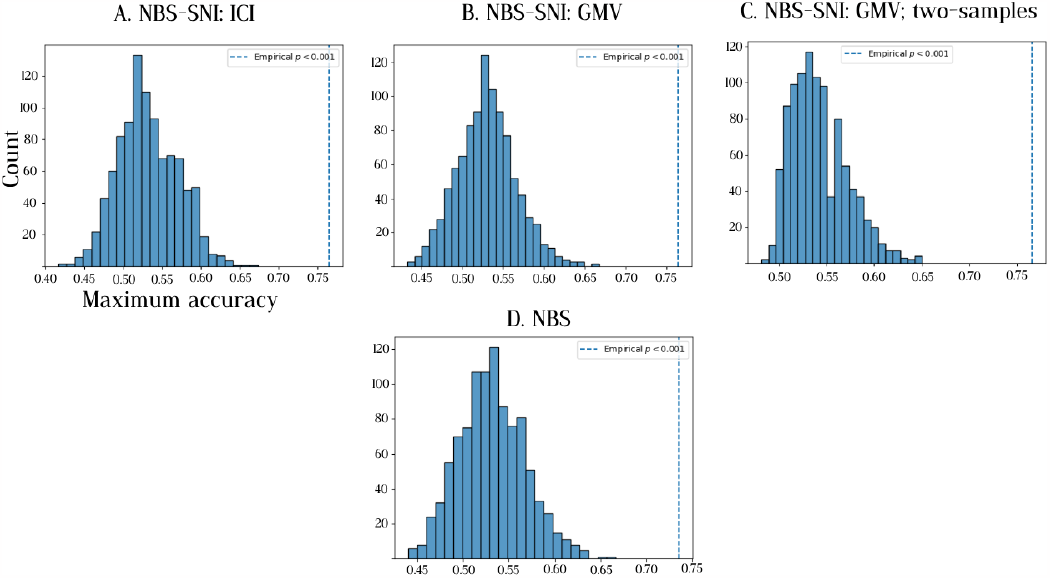
Distribution of maximum accuracy when subjects are randomly assigned to a group. A total of 1000 permutations were performed. The vertical dashed line represents the empirical prediction accuracy *μ*_performance_ obtained with the appropriate group memberships. The p-values are calculated as 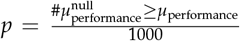 . All the prediction performances were deemed significant under this permutation test. **A** Null distribution and empirical prediction accuracy of 76.2% obtained with NBS-SNI, using the anatomical measure of intrinsic curvature index (ICI) as a node property. **B** Null distribution and empirical prediction accuracy of 77.4% obtained with NBS-SNI, using the anatomical measure of gray matter volume (GMV) as a node property. **C** Null distribution and empirical prediction accuracy of 77.6% obtained with NBS-SNI employing a two-sample t-test on the nodes, using the anatomical measure of gray matter volume (GMV) as a node property. **D** Null distribution and empirical prediction accuracy of 72.7% obtained with NBS.

## 5 CONCLUSION

In this paper, a method called NBS-SNI was introduced, which is an extension to the Network-based statistic (NBS; Zalesky et al. 2010). By being supplemented with the structural network representation, or with anatomical measures, NBS-SNI can be used to identify abnormal functional connections and perform classification in case-control studies. More precisely, the framework that we presented sets to find the most important brain regions according to a certain node property of interest (for e.g., centrality measures, anatomical measures), and probes for abnormal connectivity within or surrounding these thought-to-be important actors in the communication pathways of the brain. NBS-SNI is most well suited for cases when the contrast-to-noise ratio is relatively small in the contrast of interest. Furthermore, these connections forming the contrast should interconnect nodes that have some property which makes them stand out compared to other nodes; in that regime NBS-SNI is expected to offer a greater statistical resolution power than NBS by providing a tighter control of the family-wise error rate.

Using synthetic data, ROC curves were presented to show the regimes for which NBSSNI is expected to offer a gain in statistical resolution over NBS at identifying a contrast.

NBS-SNI was also applied on two real case-control datasets. The first one consisting of 29 individuals diagnosed with ASD and 19 healthy individuals containing both the structural and functional network representation of each individual. The second dataset comprised 75 individuals diagnosed with early psychosis and 45 healthy individuals consisting of resting-state functional networks, as well as the anatomically derived measures of intrinsic curvature index and gray matter volume.

In the fist dataset, using both network representations with NBS-SNI, a classification performance of 73% was obtained using the current flow closeness centrality of nodes, while the best performance obtained with NBS using only the functional representation was 64%. We assessed the validity of the prediction performances using permutation of the group memberships. For NBS-SNI, the prediction accuracy was deemed significant (*p* = 0.016), while the prediction accuracy of NBS was not deemed significant (*p* = 0.122).

For the second dataset, the anatomical measures of intrinsic curvature index and gray matter volume were used as a node property with our tool-box NBS-SNI. A classification performance of 77.6% was obtained with NBS-SNI using measures of gray matter volumes, while the best performance yielded by the NBS was 72.7%. All prediction performances were deemed statistically significant under the permutation test (*p <* 0.001).

We also want to emphasize the notion that NBS-SNI is a general framework, and its uses and formulation may not be limited to the ones presented in this paper. The choice of node properties to investigate is numerous. Using node properties such as centrality or anatomical measures, as it was done in this paper, seems most appropriate for large scale neuroimaging data, since the volume of the parcellated brain regions does not vary much at this scale (Simpson et al., 2013), which is essential for the soundness of the subsequent statistical analysis undertaken with NBS-SNI, which relies on a close correspondence between nodes across individuals. Another avenue for the method could be to incorporate aggregated microscale information into this framework as the node properties. Interest and recent advances in multiscale brain imaging could provide researchers with a more comprehensive understanding of the structure-function relationships (Abdeladim et al., 2019; Desjardins et al., 2019; Schirner, McIntosh, Jirsa, Deco, & Ritter, 2018; Schulz et al., 2012). Indeed, such research endeavours should involve multiscale approaches, since the relationship between structure and function is believed to be heterogeneous across the brain, due to the different physiological and molecular micro-properties of its different constituents (Khambhati, Sizemore, Betzel, & Bassett, 2018; Lariviere et al., 2019; Murphy et al., 2016; Suárez et al., 2020; van den Heuvel & Yeo, 2017). Examples of microscale information that could be incorporated in a framework like NBS-SNI include gene expression, cytoarchitechture, neuron morphology, glial cells density/type, myelin content, etc. Animals such as rats and mice are well disposed to be subject to such multiscale neuroimaging studies (Desjardins et al., 2019; Schulz et al., 2012). These kinds of datasets will become more abundant in the coming years, and we believe NBS-SNI could be a useful tool to leverage them. Alternatively, if only one brain network representation is available, node properties could also be computed directly from the functional networks.

Besides, diagnosis of certain brain conditions such as the autism spectrum disorder (ASD) can be challenging for clinical doctors, but also for the patients and their relatives. For example, a survey from 2016 in the UK shows that parents wait an average of 3.5 years between the time they approach clinical doctors to when they receive a diagnostic of autism for their children (Crane, Chester, Goddard, Henry, & Hill, 2016). Further, there are criticisms regarding the different diagnostic criteria for the different subtypes of the condition within the wider definition of the autism spectrum disorder (Tidmarsh & Volkmar, 2003). This is likely due to the overlap between the different subtypes of ASD; some are not well delineated, perhaps owing to the lack of a systematic quantification of the symptoms, brain morphology and functional activity signatures implicated in the condition. We advance that network based approaches applied in neuroscience, such as the method proposed in this work are well positioned to address these issues, and could broaden the clinical doctors’ assessments with quantitative features.

Finally, another promising avenue for patients affected by a neurodevelopmental disorder who are refractory to conventional pharmacological treatment (Lindenmayer, 2000; Siskind, McCartney, Goldschlager, & Kisely, 2016) lies in alternative therapeutic approaches, such as transcranial magnetic stimulation (TMS) (Cash et al., 2021). This approach is becoming more precise in the targeting of the brain region(s) to stimulate. We posit that the identification of abnormal functional connections between influential actors in brain networks are promising candidates for customized targeted TMS treatments.

## Limitations and recommendations

The formulation of NBS-SNI presented above only compares two properties between two groups. However, this could be extended to encapsulate more properties or more groups, or both. Multiple node or edge properties could be calculated for each of the two groups; the results of these could then be aggregated by Hotelling’s *T*^2^ test into a test statistic that is distributed along an F-distribution (Al-Labadi, Fazeli Asl, & Lim, 2022). Alternatively, data from three or more groups could be compared using ANOVA (rather than the t-tests used here), and this could then be used to select the nodes or edges for further comparison (note that this could also be used to compare more than two groups using traditional NBS). These avenues offer promise in larger scale future analyses, comparing more kinds of data from more varied datasets.

Furthermore, future analyses utilising NBS-SNI are likely to see its application in datasets with significantly larger sample sizes (e.g. the ENIGMA Consortium (Thompson et al., 2020), where studies contain thousands of participants). NBS-SNI is likely to offer statistical improvement over the canonical NBS in these datasets if neuroimaging data from multiple streams are used. However, one limitation of the present work is that the utility of NBS-SNI in large datasets has not been described. Due to the computational burden of generative network modelling (*not* the analysis of empirical data), we generated and evaluated the performance of NBS-SNI on 280 simulated networks. We therefore restricted our analyses of empirical datasets to those with a similar number of participants (given that some ground truth metrics had been established in this regime). We note that the performance of NBSSNI worsened as the CNR and sample size increased (when compared to canonical NBS; Figures 6-8); further quantification of these effects would require large scale ground-truth simulation studies. As such, we believe that NBS-SNI is best suited to research paradigms where sample size is small (*∼* 100 *−* 200 participants), CNR is relatively low, and there are complementary streams of neuroimaging data available. Although such paradigms remain common today, future usage of NBS-SNI in other regimes likely requires further modelling to validate its sensitivity and specificity.

It is important to acknowledge that it is difficult to provide definitive guidelines for the use of NBS-SNI at present. Whilst we present an integrated framework for analyzing both nodes and edges, the choice of precisely how these properties should be analyzed remains with the investigator. These choices are present at different stages in the analysis pipeline: (i) when deciding on sources of information from which to generate node and edge properties (e.g. fMRI vs diffusion MRI vs structural MRI vs alternative imaging modalities); (ii) when deciding which node or edge properties to use; (iii) when deciding what statistical thresholds to use to stratify significant nodes and edges; and (iv) when deciding which nodes and edges to retain (the parameter *d*).

Although no gold standard currently exists, some suggestions can be made regarding the options available in these domains. First, here we present examples using structural connectivity and anatomical measurements to define node properties. However, it would be reasonable to use metrics derived from alternative imaging modalities instead (e.g. as mentioned above). This provides practical benefits (given the cost and difficulty in acquiring tractographic results) and allows for the utilization of other streams of data which may be relevant to the condition under investigation. Second, the choice of which node or edge property to analyze may also be somewhat arbitrary. At this stage, we suggest that investigators consider the robustness of the imaging modality being used and the property being extracted from it. This should also be based on the extant literature regarding the condition under consideration, and multiple comparisons corrections should be used if multiple node/edge properties are analyzed. Crucially, if one input matrix comes from a noisier source, NBS-SNI may end up losing statistical power. Third, as in much of the work that utilizes NBS and similar approaches, the choice of threshold may increase the likelihood of type I or II error (if the thresholds are too low or high, respectively). Since the original conceptualization of NBS, new techniques have been developed to allow for threshold-free analyses (e.g. Baggio et al., 2018). NBS-SNI could be augmented by these techniques in future work, but at present we suggest that investigators utilize thresholds derived from the effect size expected and the sample size available. Finally, we suggest utilizing the parameter *d* = 1 at present, which balances computational demands (given that computing shortest paths is not required) and exploration of subnetworks.

## ACKNOWLEDGMENTS

FN and AA acknowledge financial support from the Sentinelle Nord initiative of the Canada First Research Excellence Fund and from the Natural Sciences and Engineering Research Council of Canada (project 2019-05183).

## REFERENCES

Abdeladim, L., Matho, K. S., Clavreul, S., Mahou, P., Sintes, J.-M., Solinas, X., … others (2019). Multicolor multiscale brain imaging with chromatic multiphoton serial microscopy. Nat. Commun., 10, 1662. doi: 10.1038/s41467-019-09552-9

Akarca, D., Vértes, P. E., Bullmore, E. T., & Astle, D. E. (2021). A generative network model of neurodevelopmental diversity in structural brain organization. Nat. Commun., 12, 4216. doi: 10.1038/s41467-021-24430-z

Al-Labadi, L., Fazeli Asl, F., & Lim, K. (2022). On Bayesian Hotelling’s T2 test for the mean. Comm. Statist. Simulation Comput., 0, 1–10. doi: 10.1080/03610918.2022.2155306

Baggio, H. C., Abos, A., Segura, B., Campabadal, A., Garcia-Diaz, A., Uribe, C., … Junque, C. (2018). Statistical inference in brain graphs using threshold-free network-based statistics. Hum. Brain Mapp., 39, 2289–2302. doi: 10.1002/hbm.24007

Baum, G. L., Ciric, R., Roalf, D. R., Betzel, R. F., Moore, T. M., Shinohara, R. T., … Satterthwaite, T. D. (2017). Modular Segregation of Structural Brain Networks Supports the Development of Executive Function in Youth. Curr. Biol., 27, 1561-1572.e8. doi: 10.1016/j.cub.2017.04.051

Baum, G. L., Cui, Z., Roalf, D. R., Ciric, R., Betzel, R. F., Larsen, B., … Satterthwaite, T. D. (2020). Development of structure–function coupling in human brain networks during youth. Proc. Natl. Acad. Sci. U.S.A., 117, 771–778. doi: 10.1073/pnas.1912034117

Betzel, R. F., Avena-Koenigsberger, A., Goñi, J., He, Y., de Reus, M. A., Griffa, A., … Sporns, O. (2016). Generative models of the human connectome. NeuroImage, 124, 1054–1064. doi:10.1016/j.neuroimage.2015.09.041

Brandes, U., Borgatti, S. P., & Freeman, L. C. (2016). Maintaining the duality of closeness and betweenness centrality. Soc. Networks, 44, 153–159. doi: 10.1016/j.socnet.2015.08.003

Cao, R., Wang, X., Gao, Y., Yuan Gao, Gao, Y., Li, T., … Xiang, J. (2020). Abnormal Anatomical Rich-Club Organization and Structural-Functional Coupling in Mild Cognitive Impairment and Alzheimer’s Disease. Front. Neurol., 11, 53–53. doi: 10.3389/fneur.2020.00053

Cash, R. F. H., Weigand, A., Zalesky, A., Siddiqi, S. H., Downar, J., Fitzgerald, P. B., & Fox, M. D. (2021). Using Brain Imaging to Improve Spatial Targeting of Transcranial Magnetic Stimulation for Depression. Biol. Psychiatry, 90, 689–700. doi: 10.1016/j.biopsych.2020.05.033

Cocchi, L., Harding, I. H., Lord, A., Pantelis, C., Yucel, M., & Zalesky, A. (2014). Disruption of structure–function coupling in the schizophrenia connectome. NeuroImage Clin., 4, 779–787. doi: 10.1016/j.nicl.2014.05.004

Crane, L., Chester, J. W., Goddard, L., Henry, L. A., & Hill, E. (2016). Experiences of autism diagnosis: A survey of over 1000 parents in the United Kingdom. Autism, 20, 153–162. doi: 10.1177/1362361315573636

Dekhil, O., Ali, M., El-Nakieb, Y., Shalaby, A., Soliman, A., Switala, A., … others (2019). A personalized autism diagnosis cad system using a fusion of structural mri and resting-state functional mri data. Front. Psychiatry, 10, 392. doi: 10.3389/fpsyt.2019.00392

Demirci, N., & Holland, M. A. (2022). Cortical thickness systematically varies with curvature and depth in healthy human brains. Hum. Brain Mapp., 43, 2064–2084. doi: 10.1002/hbm.25776

Desjardins, M., Kiliç, K., Thunemann, M., Mateo, C., Holland, D., Ferri, C. G. L., … Devor, A. (2019). Awake Mouse Imaging: From Two-Photon Microscopy to Blood Oxygen Level–Dependent Functional Magnetic Resonance Imaging. Biol. Psychiatry Cogn. Neurosci. Neuroimaging, 4, 533–542. doi: 10.1016/j.bpsc.2018.12.002

Gur, R. E., Cowell, P. E., Latshaw, A., Turetsky, B. I., Grossman, R. I., Arnold, S. E., … Gur, R. C. (2000). Reduced dorsal and orbital prefrontal gray matter volumes in schizophrenia. Arch. Gen. Psychiatry, 57, 761–768. doi: 10.1001/archpsyc.57.8.761

Gur, R. E., Turetsky, B. I., Bilker, W. B., & Gur, R. C. (1999). Reduced gray matter volume in schizophrenia. Arch. Gen. Psychiatry, 56, 905–911. doi: 10.1001/archpsyc.56.10.905

Honey, C. J., Thivierge, J.-P., & Sporns, O. (2010). Can structure predict function in the human brain? NeuroImage, 52, 766–776. doi: 10.1016/j.neuroimage.2010.01.071

Hulshoff Pol, H. E., Schnack, H. G., Bertens, M. G., van Haren, N. E., van der Tweel, I., Staal, W. G., … Kahn, R. S. (2002). Volume changes in gray matter in patients with schizophrenia. Am. J. Psychiatry, 159, 244–250. doi: 10.1176/appi.ajp.159.2.244

Khambhati, A. N., Sizemore, A. E., Betzel, R. F., & Bassett, D. S. (2018). Modeling and interpreting mesoscale network dynamics. NeuroImage, 180, 337–349. doi: 10.1016/j.neuroimage.2017.06.029

Lariviere, S., Vos de Wael, R., Paquola, C., Hong, S.-J., Mišić, B., Bernasconi, N., … Bernhardt, B. C. (2019). Microstructure-informed connectomics: enriching large-scale descriptions of healthy and diseased brains. Brain Connect., 9. doi: 10.1089/brain.2018.0587

Lindenmayer, J.-P. (2000). Treatment refractory schizophrenia. Psychiatr. Q., 71, 373–384. doi: 10.1023/A:1004640408501

Murphy, A. C., Gu, S., Khambhati, A. N., Wymbs, N. F., Grafton, S. T., Satterthwaite, T. D., & Bassett, D. S. (2016). Explicitly linking regional activation and function connectivity: Community structure of weighted networks with continuous annotation. arXiv, 1611.07962. doi: 10.48550/arXiv.1611.07962

Newman, M. E. J. (2018). Networks. Oxford University Press.

Park, H.-J., & Friston, K. (2013). Structural and Functional Brain Networks: From Connections to Cognition. Science, 342, 1238411. doi: 10.1126/science.1238411

Pascual-Belda, A., Díaz-Parra, A., & Moratal, D. (2018). Evaluating Functional Connectivity Alterations in Autism Spectrum Disorder Using Network-Based Statistics. Diagnostics, 8, 51. doi: 10.3390/diagnostics8030051

Pedregosa, F., Varoquaux, G., Gramfort, A., Michel, V., Thirion, B., Grisel, O., … Duchesnay, É. (2011). Scikit-learn: Machine Learning in Python. J. Mach. Learn. Res., 12, 2825–2830. Retrieved from https://www.jmlr.org/papers/v12/pedregosa11a.html

Pessoa, L. (2014). Understanding brain networks and brain organization. Phys. Life Rev., 11, 400–435. doi: 10.1016/j.plrev.2014.03.005

Power, J. D., Cohen, A. L., Nelson, S. M., Wig, G. S., Barnes, K. A., Church, J. A., … Petersen, S. E. (2011). Functional network organization of the human brain. Neuron, 72, 665–78. doi: 10.1016/j.neuron.2011.09.006

Roberts, G., Lord, A., Frankland, A., Wright, A., Lau, P., Levy, F., … Breakspear, M. (2017). Functional Dysconnection of the Inferior Frontal Gyrus in Young People With Bipolar Disorder or at Genetic High Risk. Biol. Psychiatry, 81, 718–727. doi: 10.1016/j.biopsych.2016.08.018

Rudie, J. D., Brown, J. A., Beck-Pancer, D., Hernandez, L. M., Dennis, E. L., Thompson, P. M., … Dapretto, M. (2013). Altered functional and structural brain network organization in autism. NeuroImage Clin., 2, 79–94. doi: 10.1016/j.nicl.2012.11.006

Schaefer, A., Kong, R., Gordon, E. M., Laumann, T. O., Zuo, X.-N., Holmes, A. J., … Yeo, B. T. (2018). Local-global parcellation of the human cerebral cortex from intrinsic functional connectivity mri. Cereb. Cortex, 28, 3095–3114. doi: 10.1093/cer-cor/bhx179

Schirner, M., McIntosh, A. R., Jirsa, V., Deco, G., & Ritter, P. (2018). Inferring multi-scale neural mechanisms with brain network modelling. eLife, 7, e28927. doi: 10.7554/eLife.28927

Schulz, K., Sydekum, E., Krueppel, R., Engelbrecht, C. J., Schlegel, F., Schröter, A., … Helmchen, F. (2012). Simultaneous BOLD fMRI and fiber-optic calcium recording in rat neocortex. Nat. Methods, 9, 597–602. doi: 10.1038/nmeth.2013

Serin, E., Zalesky, A., Matory, A., Walter, H., & Kruschwitz, J. D. (2021). NBS-Predict: A prediction-based extension of the network-based statistic. NeuroImage, 244, 118625. doi: 10.1016/j.neuroimage.2021.118625

Silva, F. N., & da F. Costa, L. (2013). Local Dimension of Complex Networks. arXiv, 1209.2476. doi: 10.48550/arXiv.1209.2476

Simpson, S., Lyday, R., Hayasaka, S., Marsh, A., & Laurienti, P. (2013). A permutation testing framework to compare groups of brain networks. Front. Comput. Neurosci., 7, 171. doi: 10.3389/fncom.2013.00171

Siskind, D., McCartney, L., Goldschlager, R., & Kisely, S. (2016). Clozapine v. first- and second-generation antipsychotics in treatment-refractory schizophrenia: systematic review and meta-analysis. Br. J. Psychiatry, 209, 385–392. doi: 10.1192/bjp.bp.115.177261

Sporns, O., & Kötter, R. (2004). Motifs in Brain Networks. PLOS Biol., 2, e369. doi: 10.1371/journal.pbio.0020369

Stephenson, K., & Zelen, M. (1989). Rethinking centrality: Methods and examples. Soc. Networks, 11, 1–37. doi: 10.1016/0378-8733(89)90016-6

Suárez, L. E., Markello, R. D., Betzel, R. F., & Misic, B. (2020). Linking Structure and Function in Macroscale Brain Networks. Trends Cogn. Sci., 24, 302–315. doi: 10.1016/j.tics.2020.01.008

Tay, J., Düring, M., van Leijsen, E. M. C., Bergkamp, M. I., Norris, D. G., de Leeuw, F.-E., … Tuladhar, A. M. (2023). Network structure-function coupling and neurocognition in cerebral small vessel disease. NeuroImage Clin., 38, 103421. doi: 10.1016/j.nicl.2023.103421

Thompson, P. M., Jahanshad, N., Ching, C. R., Salminen, L. E., Thomopoulos, S. I., Bright, J., … others (2020). Enigma and global neuroscience: A decade of large-scale studies of the brain in health and disease across more than 40 countries. Translational psychiatry, 10(1), 100. doi: 10.1038/s41398-020-0705-1

Tidmarsh, L., & Volkmar, F. R. (2003). Diagnosis and Epidemiology of Autism Spectrum Disorders. Can. J. Psychiatry, 48, 517–525. doi: 10.1177/070674370304800803

van den Heuvel, M. P., Sporns, O., Collin, G., Thomas W. Scheewe, Scheewe, T. W., Scheewe, T. W., … Kahn, R. S. (2013). Abnormal Rich Club Organization and Functional Brain Dynamics in Schizophrenia. JAMA Psychiatry, 70, 783–792. doi: 10.1001/jamapsychiatry.2013.1328

van den Heuvel, M. P., & Yeo, B. T. (2017). A spotlight on bridging microscale and macroscale human brain architecture. Neuron, 93, 1248–1251. doi: 10.1016/j.neuron.2017.02.048

Van Essen, D., & Drury, H. (1997). Structural and functional analyses of human cerebral cortex using a surface-based atlas. J. Neurosci., 17, 7079–7102. doi: 10.1523/jneurosci.17-18-07079.1997

Van Essen, D. C., Smith, S. M., Barch, D. M., Behrens, T. E., Yacoub, E., Ugurbil, K., … others (2013). The wu-minn human connectome project: an overview. NeuroImage, 80, 62–79. doi: 10.1016/j.neuroimage.2013.05.041

Vita, A., De Peri, L., Deste, G., & Sacchetti, E. (2012). Progressive loss of cortical gray matter in schizophrenia: a meta-analysis and meta-regression of longitudinal mri studies. Transl. Psychiatry, 2, e190–e190. doi: 10.1038/tp.2012.116

Wang, J., Khosrowabadi, R., Ng, K. K., Hong, Z., Chong, J. S. X., Wang, Y., … Zhou, J. (2018). Alterations in Brain Network Topology and Structural-Functional Connectome Coupling Relate to Cognitive Impairment. Front. Aging Neurosci., 10, 404–404. doi: 10.3389/fnagi.2018.00404

Wen, T., & Deng, Y. (2020). Identification of influencers in complex networks by local information dimensionality. Inf. Sci., 512, 549–562. doi: 10.1016/j.ins.2019.10.003

White, T., Andreasen, N. C., Nopoulos, P., & Magnotta, V. (2003). Gyrification abnormalities in childhood-and adolescent-onset schizophrenia. Biol. Psychiatry, 54, 418–426. doi: 10.1016/S0006-3223(03)00065-9

White, T., & Hilgetag, C. C. (2011). Gyrification and neural connectivity in schizophrenia. Dev. Psychopathol., 23, 339–352. doi: 10.1017/S0954579410000842

Witvliet, D., Mulcahy, B., Mitchell, J. K., Meirovitch, Y., Berger, D. R., Wu, Y., … Zhen, M. (2021). Connectomes across development reveal principles of brain maturation. Nature, 596, 257–261. doi: 10.1038/s41586-021-03778-8

Zalesky, A., Fornito, A., & Bullmore, E. T. (2010). Network-based statistic: Identifying differences in brain networks. NeuroImage, 53, 1197–1207. doi: 10.1016/j.neuroimage.2010.06.041

Zhan, Y., Yao, H., Wang, P., Zhou, B., Zhang, Z., Guo, Y., … Liu, Y. (2016). Network-Based Statistic Show Aberrant Functional Connectivity in Alzheimer’s Disease. IEEE J. Sel. Top. Signal Process., 10, 1182–1188. doi: 10.1109/JSTSP.2016.2600298

Zhang, R., Shao, R., Xu, G., Lu, W., Zheng, W., Miao, Q., … Lin, K. (2019). Aberrant brain structural–functional connectivity coupling in euthymic bipolar disorder. Hum. Brain Mapp., 40, 3452–3463. doi: 10.1002/hbm.24608

